# Diversity and genetic structure within a Mexican maize race reveal consistent biocultural processes across geographic scales

**DOI:** 10.1101/2025.10.28.685174

**Authors:** Duhyadi Oliva-García, Bulmaro Coutiño-Estrada, Idalia C. Rojas-Barrera, Hugo Perales, Daniel Piñero, Ana Wegier, Alicia Mastretta-Yanes

## Abstract

Understanding how genetic diversity is distributed within maize races is critical for conserving maize genetic diversity. While maize races are traditionally treated as homogeneous units to be targeted by conservation, each race may encompass genetically distinct populations of landraces, shaped by environmental conditions and farmer selection. In this study, the genetic diversity and population structure of the *Olotillo* race were examined using 89,810 SNPs from 158 samples collected across local, regional, and national scales in Mexico. Additional samples from other races *Dzit-bacal*, *Tuxpeño*, and a *Mix*, group combining *Olotillo* with other varieties, brought the total to 171 individuals. Population structure analyses (ADMIXTURE, PCA) revealed that genetic clusters do not align with race identity, but instead reflect geographic origin. Inbreeding depression was evident within *Olotillo* populations, with mean *F*_IS_ = 0.746 ± 0.066. A heterozygote deficiency was observed: mean *H_e_*OB = 0.075 ± 0.020, compared to a higher expected heterozygosity (*H_e_*EX = 0.295 ± 0.002). Expected homozygosity (*H_O_*EX) of 0.705± 0.002 was observed, reflecting reduced effective genetic variation. The above is reflected in the population contraction signal (e.g., positive Tajima’s D), and no significant differences between scales in *H*_e_EX, *H_O_*EX, and nucleotide diversity (π), suggesting that similar evolutionary forces act across regions with a strong directional selection. These findings underscore the need to ascertain the genetic background among races and to recognize meaningful within-race differentiation. Conservation actions should reflect this structure, promoting targeted seed exchange to counteract genetic erosion while respecting farmer preferences. *Olotillo* exemplifies how maize diversity is structured not only by genetics, but by cultural and management systems that shape evolution *in situ*.

## Introduction

The Kunming-Montreal Global Biodiversity Framework (GBF) aims to maintain the genetic diversity within populations of wild and domesticated species to maintain their adaptive potential (CBD 2022). For crop species, like maize, this opens up the question of how genetic diversity is distributed within them in order to conserve genetic diversity at the population level where evolutionary processes take place. Maize is one of the three cereals that feed the world (Nuss & Tanumihardjo 2010) and one of the most celebrated crops for its cultural importance from North America to South America (Staller 2009, Wertz 2005). Depictions of maize tend to allude to its great morphological diversity, with the number of native maize races often used to express maize genetic diversity within a region. Here, we argued that conserving maize genetic diversity requires going beyond the race unit, as what is morphologically classified as a single race may encompass hundreds of local varieties known as landraces (Bellon et al. 2018; Palacios-Paola et al. 2022).

Due to its cross-mating system, maize (*Zea mays* L. subsp. *mays*) is characterized by concentrating ∼80% of the genetic diversity of teosintes, its wild ancestors (Hufford et al. 2012). With 59 recognized native races (Ron et al. 2006; Sánchez et al. 2000) and being the center of origin of the crop, Mexico holds the greatest maize genetic diversity (Prasanna, 2012). This diversity is still evolving in the hands of millions of small-holder farmers (Rojas-Barrera et al. 2019), who continue the process of domestication by selecting, growing and saving their own seed every cycle (Bellon et al. 2018). Thus, maize races are morphological characterizations that aim to reflect a natural classification (Wellhausen et al. 1951; Goodman & Paterniani 1969; Sánchez et al. 1993; Sánchez & Goodman 1992), but thousands of farmers can grow a single race in localities that can be hundreds or thousands of kilometers apart (Rice E. 2007). Therefore, maize races are likely conformed by several populations of landraces, with different selection pressures, isolation levels, and gene flow, including with other maize populations.

In Mexico, landraces are mostly grown in regions of smallholder “campesino” agriculture, comprising more than four million hectares planted with non-commercial varieties (Bellon et al. 2018), which nevertheless respond to the demand of non-maize producing local consumers (Bellon et al. 2021). From a cultural point of view, landraces are maintained *in situ* through the traditional knowledge systems that originate and conserve them. From an evolutionary point of view, that maize is cultivated in a large territory and under different environmental conditions, implies a great capacity for local adaptation to various conditions (Bellon & Risopoulos 2001). Indeed, farmers identify which maize landrace best suits their field conditions (Hellin et al. 2014), which translates into locally adapted maize populations that align with local cultural preferences (Bellon et al. 2003; Caballero et al. 2023). Concurrently, consumption preferences are recognized in certain varieties, such as nutritional and taste preferences (Bellon, 1996; Bellon & Hellin, 2011; Palacios-Paola et al. 2022). In addition, they are part of the traditions and customs (Ocampo-Giraldo et al. 2020), and thus emerge from a co-evolutionary process between maize and farmers (Abraham-Juárez et al. 2021). This co-evolutionary process (Bellon, 1991; Purugganan 2022) has been maintained by approximately 60 indigenous groups and non-indigenous farmers in Mexico (Arteaga et al. 2016).

Although landraces within maize races are recognized for their cultural and biological importance, how genetic diversity is distributed among them remains understudied. This distribution is shaped by seed exchange and pollen movement; maize pollen typically travels up to 30 meters (Bøhn et al. 2016) or more, while seed exchange enables broader gene flow (Mercer & Wainwright 2008). The relative impact of these processes depends on field size, geography, and local practices, underscoring the need for high-resolution genetic data (Mercer & Wainwright 2008). Despite this, landraces have been relatively poorly studied using genomic data relative to other aspects of maize domestication. For instance, studies using several individuals per race were done using microsatellites (Pineda-Hidalgo et al., 2013) instead of genomic data. This may underestimate diversity because genomic data (i.e. thousands of SNPs) provides more precise estimates of population-level diversity, higher power to identify groups in clustering methods, and the ability to consider local adaptation (Zimmerman et al 2020). Similarly, a study using genomic data focused on few (1-16) individuals per race, most of them from the same locality (Arteaga et al. 2016), thus failing to represent population-level differentiation. Therefore, given that genetic diversity (Arteaga et al. 2016) and agronomic variation (Torres-Morales et al. 2022) within each race have been little explored at the population level, the genetic diversity of Mexico’s native maize may have been underestimated.

This paper examines genetic diversity and population structure within a single maize race across different geographic scales. Seed exchange varies between local, regional, and national levels (Brush & Perales 2007; Dyer & Taylor 2008), with varying degrees of gene flow and environmental variation expected at each scale. Therefore, understanding the genetic diversity and population structure within a single maize race across different geographic scales is important. Local exchange occurs within the same community or municipality, typically among neighbors or extended family members. This type of exchange is widely documented and is the most frequent (Badstue et al. 2006; Dyer & López-Feldman 2013; Llamas-Guzmán et al. 2022). Regional exchange occurs between communities in neighboring municipalities and is less common than local exchanges (Brush & Perales 2007). Although less frequent, regional exchange has been documented when farmers seek seeds that differ from those available locally (Badstue et al. 2006). National exchanges, which are the least frequent, involve exchanges between different states (Brush & Perales 2007). National exchange can be linked to current long-distance trips (Badstue et al. 2006) but signatures of historical human migration over the last two centuries, when the Mexican population grew and expanded after the Mexican Revolution, may also be present.

Hypothesizing that genetic diversity within a race differs at local, regional, and national levels, potentially due to management practices, geographic distance, and environmental variation, the Olotillo race is a suitable system to test this hypothesis. In Mexico, this race has high cultural importance for the families that cultivate it within a wide range of temperatures and rainfall (**Fig. 1**). Despite this, *Olotillo* is planted in smaller areas than what it used to be, and fewer farmers are involved in growing it, as it has been largely displaced by hybrid maize since the last century (Bellon & Risopoulos 2001).

**Fig. 1.**
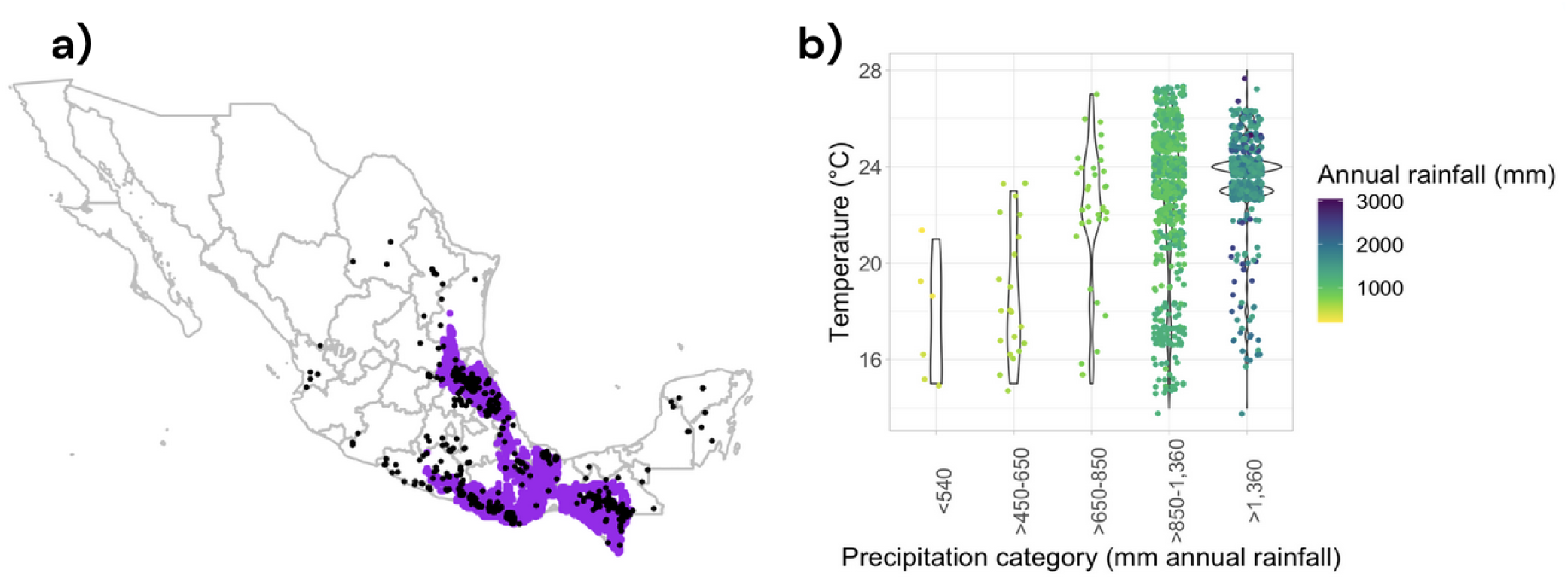
Distribution and environmental conditions where *Olotillo* is grown. **a)** Purple color shows the climatic interpolation of *Olotillo* distribution, restricted to the main altitude where it is grown, according to Perales & Golicher (2014). Black points are the sampling points of *Olotillo* from the Global Maize Project (GMP: CONABIO, 2011). **b)** Range of temperature and precipitation of *Olotillo’*s sampling points from the GMP. Temperature (°C) refers to the mean annual temperature for the 2000 period, and precipitation (mm) to the mean annual rainfall for the same period, which were obtained from Cuervo-Robayo et al (2020) for each of the sampling localities of the GMP. Precipitation range categories were defined based on Ruiz et al. (2008). Data was downloaded from the “Explorador de Condiciones Climáticas de Agrobiodiversidad” of the Sistema de Información de Agrobiodiversidad (app-siagro.conabio.gob.mx).

The name *Olotillo* derives from the Nahuatl *olotl,* (heart in Spanish) in combination with the diminutive *illo*, which means “maize with small cob”, in reference to the thin cobs of this race (Wellhausen et al. 1951). The eight rows and cob flexibility are the other important distinguishing features of this race. It is worth noting the influence of *Olotillo* on other races: *Tuxpeño*, *Vandeño*, *Chalqueño*, *Celaya*, and *Cónico Norteño* (Wellhausen et al. 1951).

*Olotillo* is widely grown, though not abundant, in the Central Depression of Chiapas state, Mexico (Bellon,1991), one of the states with the greatest genetic diversity of maize (Perales & Hernández, 2005). Its main distribution center is the upper Grijalva River, with presence and influence towards the coastal regions and slopes of the Pacific slope in Oaxaca and Guerrero, as well as the coastal areas, foothills, and slopes of the Gulf of Mexico, from the north of Oaxaca to Veracruz, Puebla, Hidalgo and San Luis Potosí states (CONABIO, 2020). *Olotillo* is grown mostly in lowlands, tropical dry forest environments, and pine-oak forests. It also has been reported to grow at 2,000 masl (Callejas-Chavero et al. 2019) but these are likely misidentifications (Brush & Perales 2007). Regarding ecological features, *Olotillo* has a long cycle, drought resistance, weed resistance, and the ability to grow in poor soils. However, it retains a tall height, is prone to lodging due to its weak stalk, and has limited compatibility for growth when intercropped with squash (Bellon,1991). In terms of management, it is characterized by needing less maintenance than other races, because, in the words of the farmers “it is resistant and not delicate” (“*aguantador”* and “*no delicado*” in Spanish). It has a low yield relative to its weight, poor storage qualities, and little presence in the market because it is used mainly for self-consumption. It is sought for its good taste and in traditional markets is sold by volume, which offsets its low yield (Bellon,1991). Despite the former disadvantages, *Olotillo* is very appreciated gastronomically, and it is an important component of local maize consumption, both as “elote”, “tortilla”, “pozole”, and other dishes (farmers personal communications during fieldwork). Women especially appreciate its culinary properties (farmers personal communications during fieldwork).

*Olotillo* landraces were sampled at the local, regional, and national levels, covering an important part of its known distribution and emphasizing Chiapas state, where they are widely cultivated. By describing the distribution of genetic diversity at this level of detail within a single race, we aim contributing to a broader understanding of how landrace diversity is maintained, reinforcing the need for nationwide efforts that support the farmers managing the evolutionary processes sustaining crop genetic diversity (Mastretta-Yanes et al., 2024). For this, we first seek to demonstrate that maize races are not genetically homogeneous units, and therefore require monitoring and conservation efforts at the population or landrace level. Then, we intend to inform with genetic data both *in situ* and *ex situ* conservation strategies for maize races like *Olotillo,* which, despite their high cultural value and adaptation to diverse environments, are being cultivated less frequently.

## Material and methods

### Plant material

The maize accessions for the *Olotillo* race were sampled in Mexico from 2018 to 2019 on three different geographic scales: local (distance range: 40.9 Km), regional (distance range: 55 Km), and national. Sampling covered a wide part of the known distribution of this race and environmental conditions (**Fig. 1**). For the local level, 12 accessions were sampled: 1 from Jiquipilas and 11 from Ocozocoautla, both municipalities in the state of Chiapas. Local level collections refer to accessions that were collected in the same municipality. The Jiquipilas sample was considered local because of its proximity to Ocozocuatla. At the regional level, 8 accessions were collected from municipalities surrounding Ocozocoautla: 1 from Villaflores, 2 from Cintalapa, 1 from Suchiapa, 1 from El Parral, 1 from Chiapa de Corzo and 2 from Comitán de Domínguez. At the national level, samples were from 2 communities in the state of Oaxaca (collected by Flavio Aragón Cuevas), 11 from the state of Guerrero (by Noel Gomez Montiel), and 4 from the state of Nayarit (by Víctor Vidal Martínez). To complement the geographic reach of our sampling, *Olotillo* accessions from CIMMYT (International Maize and Wheat Improvement Center, Texcoco, Mexico) were also incorporated from the states of Hidalgo, San Luis Potosí and Veracruz (**Fig. 2**). Our sampling and CIMMYT’s accessions are compatible because both focus on point localities within the sampled states. A total of 49 accessions were collected, from which 171 samples (individual seeds) were obtained (**Table 1)**. Location details are available in **Table S1.** Additionally, two more races were included, *Tuxpeño* and *Dzit-Bacal*. The first is to compare genetic diversity structure with a race that is now displacing *Olotillo* and is now being planted at higher proportion due to higher yields (López-Morales et al. 2017). *Tuxpeño* is considered an intermediate race between *Olotillo* and *Tepecintle* (Sierra-Macías et al. 2016) and reported as one of the most productive worldwide for 1974 (Bellon & Brush 1994). The *Dzit-bacal* race was included to compare diversity structure against a sister race (Wellhausen et al. 1951). For the *Tuxpeño* race, two accessions from Campeche were sampled. For the *Dzit*-*Bacal* race, three accessions were sampled coming from Campeche (2 samples) and Quintana Roo (1 sample) states.

**Fig. 2.**
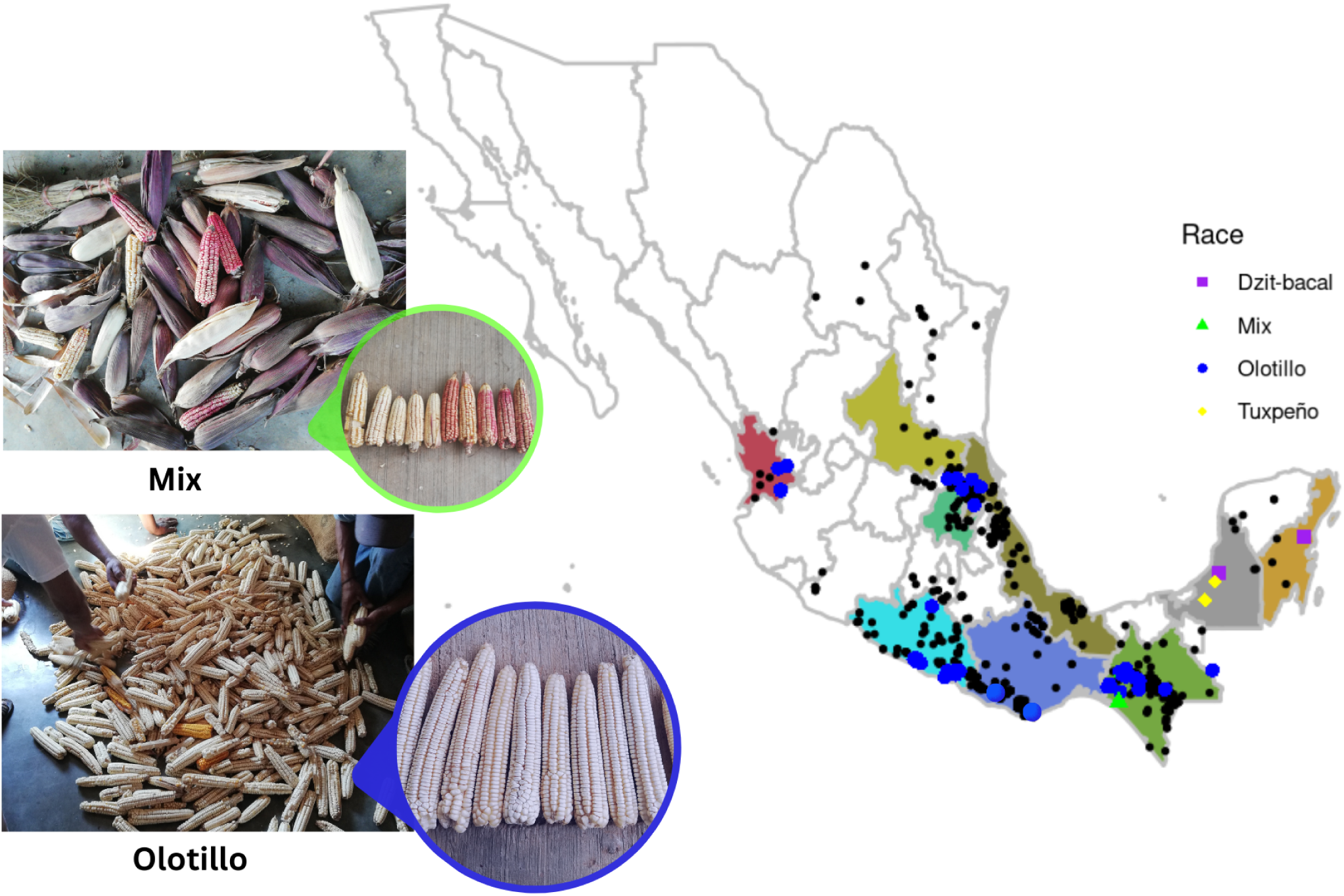
Sampled locations for this study. Black points: *Olotillo* accessions according to the Global Maize Project (CONABIO, 2011). Colored dots: Localities sampled for this study are colored by race. The photos were taken during sampling. The image of what we call *Mix* corresponds to an accessionmade up of a mixture of the following landraces of maize: *Cuarenta y dos*, *Cuarentan*o, *Olotillo morado*, *Diente de venado,* and *Jarocho*, based on the farmer’s identification from Tonalá, Chiapas. The photo *Olotillo, white and yellow* is from Suchiapa, Chiapas. Notice the thin cob of *Olotillo*.

The key traits used to identify the *Olotillo* race in the field included relatively long ears with predominantly eight rows and a thin, flexible cob (Welhausen et al. 1951; van Heerwaarden et al. 2009), and the acknowledgment of local farmers. The *Olotillo* samples that were included in this study were of white color. Additionally to the *Dzit-bacal* and *Tuxpeño* races, a mixture (*Mix*) of different varieties (*Olotillo*, *Rocamey*, and *Veracruzano;* CHIS_L38, **Table S1**) was included to evaluate the effect of mixing different maize varieties, a common practice among campesinos.

Sampling was performed in small villages, usually with less than 20,000 inhabitants. In each locality, one to three campesinos family farms --traditional farmers usually with less than 5 ha of land (Bellon et al. 2018)-- were visited. Farmers still cultivating *Olotillo* were identified and invited to participate in interviews. The objectives of the project and the motivation for studying *Olotillo* were explained to them, and upon agreement, 100 ears from the most recent growing season (July 2018) were collected from each farmer, forming part of the “*Olotillo* Evolutionary Breeding Project” accessions set. This sample size was chosen to minimize problems associated with inbreeding (Crossa et al. 1994; Mercer & Perales 2019). In some cases, fewer than 100 ears were obtained, or pre-shelled seeds were provided instead (see **Table S1**). Recognizing that 100 ears may represent a considerable portion of the harvest for traditional farmers, their contribution was compensated with an amount equivalent to the local market value of the seed.

### DNA extraction and sequencing

Four individual maize plants per accession were sequenced, for which eight seeds were first germinated in an environmental chamber (CONVIRON) at 25°C in 12 h of darkness and light. Extraction was performed on four of the eight plants per accession in 7-day-old seedlings, and the amount of tissue was one leaf (2.5 cm length). The DNesay Plant Mini Kit (QUIAGEN, Inc., Germantown, MD, USA) was used to extract DNA. The tissue grinding was performed by placing the fresh tissue in a 2 ml tube and adding liquid nitrogen. The grinding was then carried out within the same tube using a plastic pestle. DNA quality was evaluated using 1% agarose gel electrophoresis. DNA concentration was quantified using fluorometry, Qubit 3.0 (Thermo Fisher Scientific Inc., Waltham, MA, USA) with a Qubit dsDNA BR Assay Kit. DNA samples from each individual (germinated plant) were sent to the University of Wisconsin Biotechnology Center (Madison, WI, USA) following a GBS protocol (Wang et al. 2020) using the ApeKI enzyme (Rojas-Barrera et al. 2019). Two plates, each with 95 samples plus a blank, were sequenced. This study used 171 samples (individual maize plants), including 158 for *Olotillo*, 4 for *Tuxpeño*, 6 for *Dzit-bacal*, and 3 for the *Mix*.

### Bioinformatics workflow

#### Variant calling (SNPs), and data quality assessment

The quality readings were carried out with FastQC (https://www.bioinformatics.babraham.ac.uk/projects/fastqc/). Fastq files were demultiplexed with GBSx v1.3 (Herten et al. 2015). Reads alignment was performed with Nextgenmap 0.5.3 (Sedlazeck et al. 2013) against the B73 assembly version: Zm-B73-REFERENCE-NAM-5.0 (https://download.maizegdb.org/Zm-B73-REFERENCE-NAM-5.0). Subsequently, aligned reads were converted into binary files using Samtools 1.5 (Li et al. 2009). Variant calling was performed with HaplotypeCaller and genotype merging was performed with GenotypeGVCFs, both tools of the Genome Analysis Toolkit, GATK 3.8.0 (McKenna et al. 2010). SNP filtering parameters were determined based on the distribution of SNP calling metrics, including quality, variant loss, mean depth, and the lowest frequency alleles per site and individual, computed using VCFtools v0.1.16 (Danecek et al. 2011), see Supplementary Methods (**Table S2, Fig. S1-Fig. S7**) for details on parameter selection. Correctly choosing the filtering parameters is directly related to the quality of the values in common population genetics statistics (Hemstrom et al. 2024). The filtering arguments were **--**maf 0.05 **--**max-alleles 2 **--**max-missing 0.80 **--**min-meanDP 0.5 **--**max-meanDP 4 **--**minDP 2 **--**maxDP 4. A total of 89,810 SNPs out of 4,959,703 SNPs were retained at the end of the processing.

#### Analysis of population structure

Two unsupervised analyses, PCA (Price et al. 2006) and ADMIXTURE (Alexander et al. 2009), were applied to explore population genetic structure. PCA was conducted using the SNPRelate (Zheng et al. 2012), R package (R Core Group 2024), for which only biallelic SNPs were extracted at the moment of conversion, thus thePCA was performed using 76,543 biallelic SNPs. To characterize the genetic structure with cluster composition at the individual level with the Bayesian approach, ADMIXTURE v1.3.0 was used which was run 10 times for each K, obtaining the CVE for each run (see Supplementary Methods for CVE’s plot).

#### Genetic diversity

Using the set of 89,810 SNPs, genetic diversity values were calculated by race (**Table 2**) and by geographical scale (**Table 3**). First, the --het command of VCFtools was employed to estimate the number of observed homozygous sites (OBs(_HOM_)), the number of expected homozygous sites (EXs(_HOM_)), and the inbreeding coefficient (*F*_IS_) for each sample. Following the approach applied by Armstrong et al. (2024), the number of heterozygous sites was derived from these values as follows: observed heterozygous sites (OBs_(HET)_) = Number sites - OBs_(HOM)_, expected heterozygous sites expected (EXs_(HET)_) = Number sites - EXs_(HOM)_, observed heterozygosity (*H_e_OB*) = OBs_(HET)_ / Number sites, and expected heterozygosity (*H_e_*EX)= EXs_(HET)_ / Number sites. Finally, expected homozygosity was obtained from (*H_O_*EX) = 1 - *H*_e_EX. Mean and standard deviation of genetic diversity summary statistics for each accession are shown in **Table S3**. To obtain the nucleotide diversity (π), the value was estimated from 10 kb (10 000 pb) windows across the genome, in VCFtools with --window-pi 10000. For Tajima’s D, the calculation was also done in a window of 10 kb, --TajimaD 10000. Estimating nucleotide diversity from GBS data using VCFtools may underestimate genetic diversity due to missing genotypes, either from missing or intrinsic characteristics of the GBS approach (Korunes & Samuk, 2021). Nonetheless, this bias is systematic and thus expected to affect all samples uniformly. Statistical analyses and plots were made with the R (R Core Team, 2025) package ggstatsplot version 0.13.1 (Patil, 2021).

## Results

After data processing, a total of 89,810 SNPs (filtered data set) were obtained from all accessions (n = 49) and samples (n = 171). Of these, 158 corresponded to *Olotillo*, 4 to *Tuxpeño*, 6 to *Dzit-Bacal,* and 3 to *Mix* (**Table 1**). Population structure at the state and different geographical scales, analysed with 76,543 biallelic SNPs for the other races, including *Mix*, was supported by the first two principal components, which explained 1.58% (PC1) and 0.97% (PC2) of the total variability (**Fig. 3**).

**Fig. 3.**
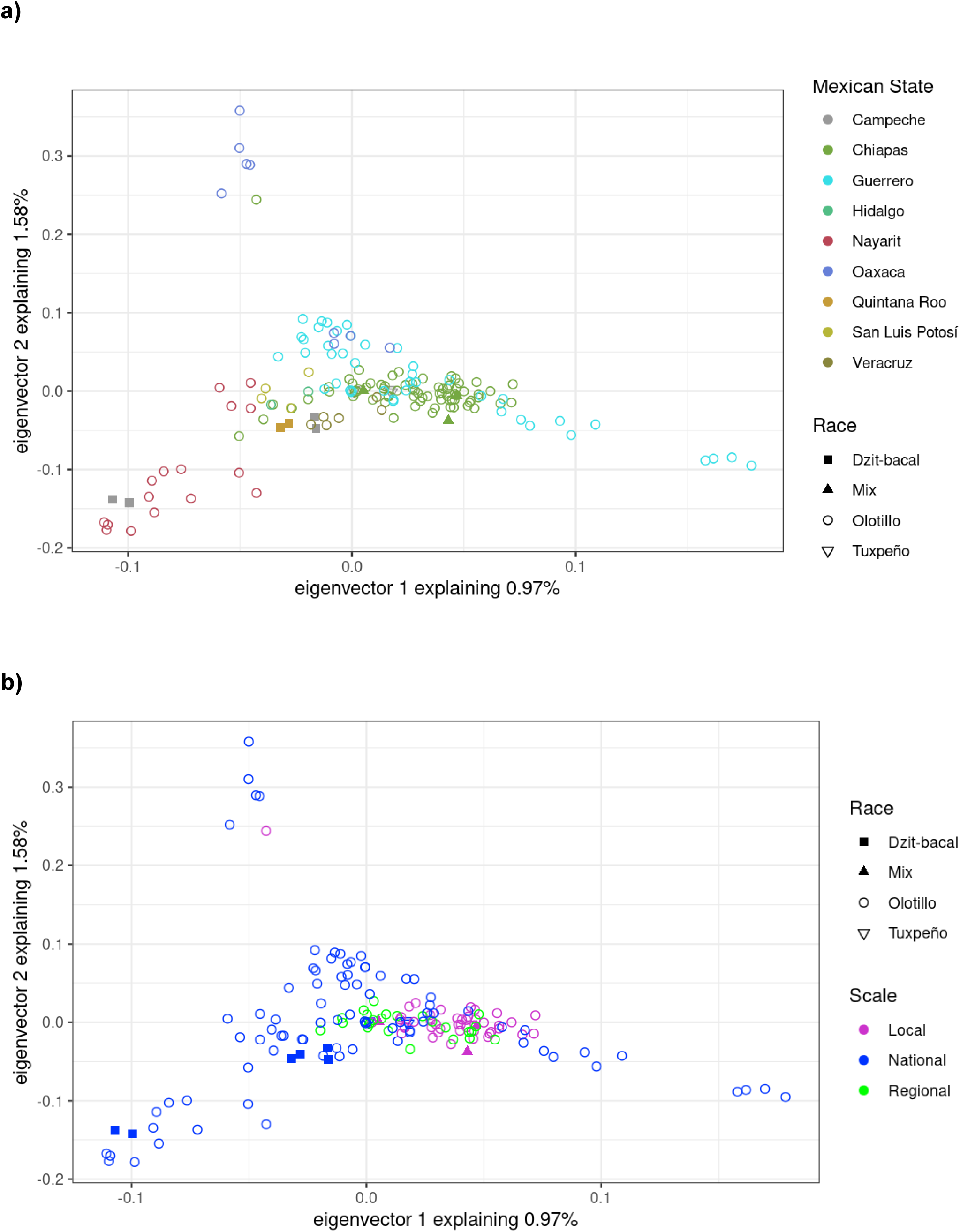
Principal component analysis (PCA) using 76, 543 SNPs. The first two components are shown **a)** By state: state color points as in Fig. 1 **b)** By scale: local, regional, and national.

The results of the ADMIXTURE analysis, including all samples together (**Fig. 4**), show how the genetic differentiation increases when evaluating *K* values from 2 to 4 (number of genetic clusters). For *K* = 2, individuals are mainly grouped into two clusters, suggesting an initial differentiation between two large and predominant genetic groups in all races. The same pattern holds for *K* = 3 and *K* = 4, with all genetic clusters present in all races, but in different proportions. *K* = 2 had the smallest value for the cross validation error (**Fig. S8**). The Admixture clusters do not correspond to races, which would have been expected if races were genetic units characterized by shared ancestry and low gene flow with other races. Instead, structure is observed both within and between races.

**Fig. 4.**
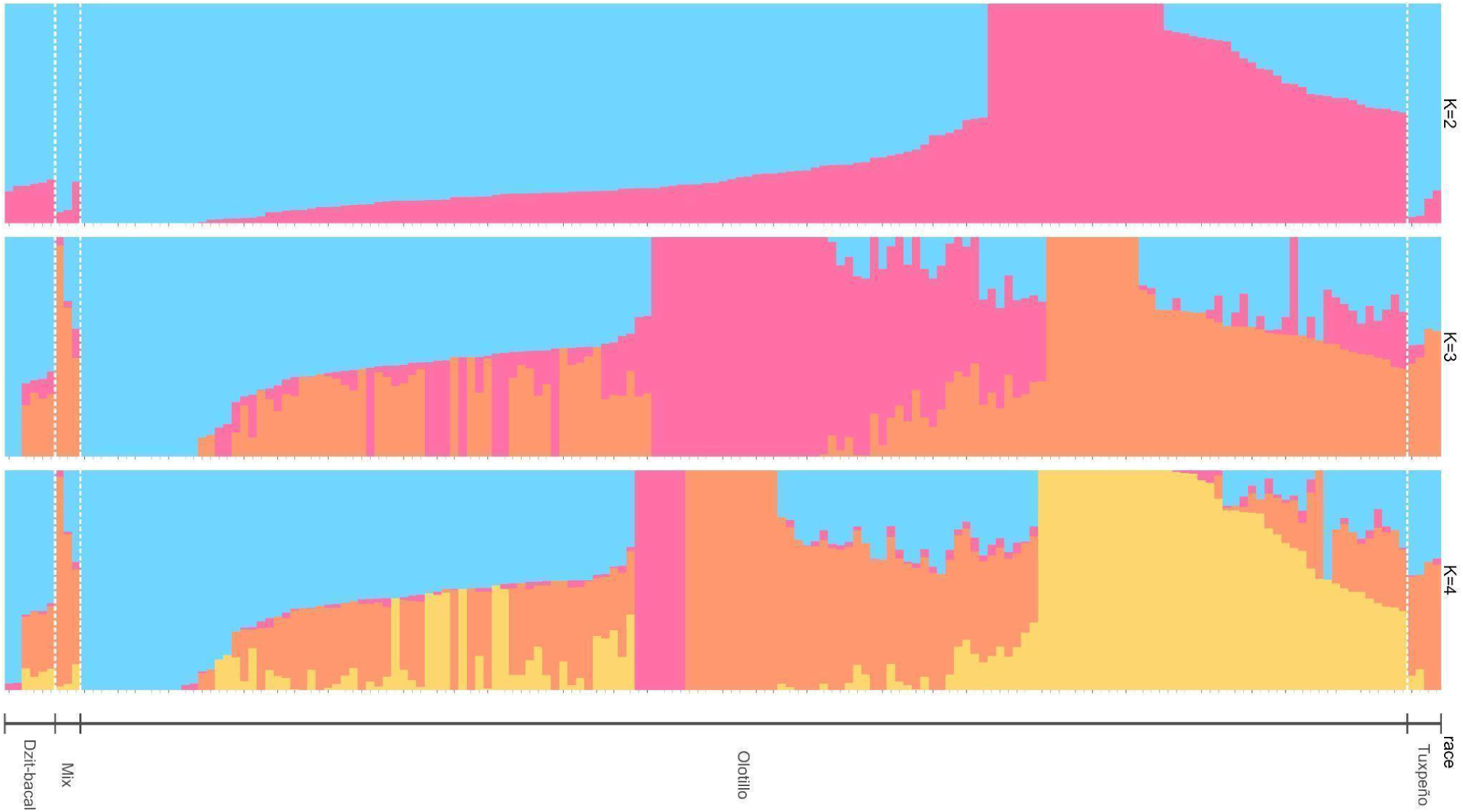
Genetic structure for all samples. Population structure analysis for three values of *K* for *Dzit-bacal*, *Olotillo,* and *Tuxpeño* races, as well as for an accession of mixed races (*Mix*). Each vertical bar represents the membership probability (0-1.0) for each sample (an individual), and the colors inside the bars indicate the likelihood of the genome assigned to each genetic cluster. Samples are ordered by cluster membership within each race. *K* = 2 had the lowest CVE value.

In **Fig. 5**, an ADMIXTURE analysis only for the *Olotillo* race is shown, with samples grouped by state and scales: local, regional, and national. Two genetic clusters (*K*= 2) were observed for *Olotillo*, suggesting a geographic differentiation within the *Olotillo*, with one cluster being the most predominant at the local and regional scales, sampled in Chiapas. When *K* = 3, the third cluster differentiates the samples from Guerrero, although it is also present in other states and scales. This increase in genetic resolution allows us to identify finer differences between *Olotillo* populations. In *K* = 4, the fourth genetic cluster is again mostly present in the local and regional sampling within Chiapas and San Luis Potosí (SLP) at the national level. Overall, the admixture analysis within *Olotillo* shows that particular genetic clusters are more prevalent in certain States, suggesting geographic genetic differentiation even within a single race.

**Fig. 5.**
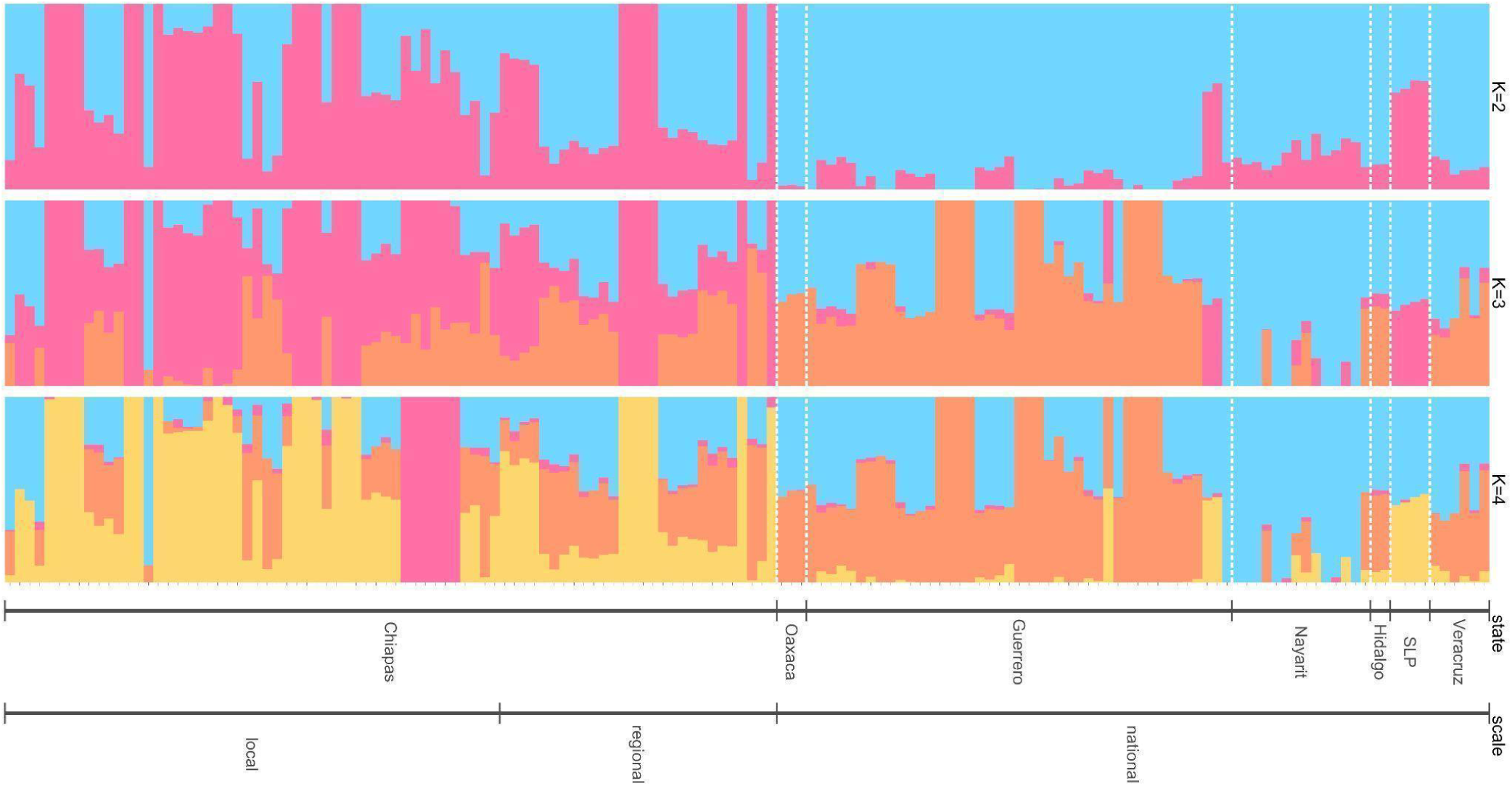
Genetic structure within *Olotillo*. Population structure analysis for three values of *K* for *Olotillo*. Each vertical bar represents the membership probability for each sample (an individual), and the colors inside the bars indicate the likelihood of the genome assigned to each cluster. Samples are grouped by scale (local, regional, and national) and then by state. *K* = 2 had the lowest CVE value.

**Table 2** presents the population genetics statistics by race. However, due to sampling size differences (*Olotillo* n=158 vs n=3,4 and 6 for the other races) it is not possible to statistically compare *Olotillo* and the other races. **Table 3** presents the population genetics statistics for *Olotillo* by geographic scale. The mean *F*_IS_ values for the *Olotillo* maize landrace showed at the Local with lowest levels of inbreeding (*F*_IS_ = 0.715), followed by National (*F*_IS_= 0.750) and finally Regional (*F*_IS_ = 0.759). The Kruskal-Wallis revealed significant differences (*p* = 0.03, *p* < 0.05) in the median for *F*_IS_, Local (*F*_IS_ = 0.740), Regional (*F*_IS_ = 0.759) and National (*F*_IS_ = 0.743). The post hoc Dunn’s Multiple Comparison confirmed that there are significant differences between Local and Regional (*p* = 0.04, *p* < 0.05), and no significant differences between Local and National (*p* = 0.06, *p* > 0.05), and Regional and National (*p* = 0.39, *p* > 0.05), **Fig. S9**. The mean *H*_e_OB with the lowest was Regional (*H*_e_OB = 0.071), followed by National (*H*_e_OB = 0.074) and finally Local (*H*_e_OB = 0.084). The Kruskal-Wallis revealed significant differences (*p* = 0.03, *p* < 0.05) in median between Local (*H*_e_OB = 0.077), Regional (*H*_e_OB = 0.071) and National (*H*_e_OB = 0.076). Post hoc Dunn’s show significant differences between Local and Regional (*p* = 0.04, *p* < 0.05), and no significant differences between Local and National (*p* = 0.06, *p* > 0.05), and Regional and National (*p* = 0.39, *p* > 0.05), **Fig. S10**.

The mean and median values for *H*_e_EX were very similar between scales, therefore no significant differences were found, **Fig. S11**, same results in *H*_O_*EX*, **Fig. S12**. For the value of π, the Kruskal-Wallis test also showed significant differences (*p* = 0.03, *p* < 0.05). However, no significant differences were found in post hoc Dunn’s test, **Fig. S13**. For Tajima’s *D* significant differences were found in Kruskal-Wallis test (*p* = 0.00, *p* < 0.05) and in post hoc Dunn’s test, **Fig. S14.** Finally, no significant differences were found in the *N_s_* for Kruskal-Wallis test (*p* = 0.15, *p* < 0.05), **Fig. S15**.

## Discussion

This study provides novel insights into the genetic diversity and structure of the *Olotillo* maize race using extensive sampling and 89,810 SNPs. Unlike previous studies with limited within-race representation, our results show that there is genetic structure within the *Olotillo* race, and that this is related to geography (**Fig. 5**). Additionally, evidence of population contraction and inbreeding was identified, likely driven by traditional seed practices and reduced cultivation (Bellon et al.1991). These findings emphasize the need to reconsider race as the main unit for maize genetic diversity conservation, and inform strategies for the *in situ* conservation of native maize.

### Genetic differentiation within the Olotillo race

Previous studies on genetic diversity within maize races using genomic data have focused on several races, but including few samples per race (Arteaga et al. 2016; Caldu-Primo et al. 2017; Rocandio-Rodríguez et al. 2014). In contrast, this study focused on the *Olotillo* race with an extensive sampling across the Mexican states where it is grown, encompassing 158 samples of this race, as well as three other races. The population structure analyses show genetic structure within *Olotillo* (**Fig. 5**), but the same genetic clusters are also found for the other sampled races from the same state (**Fig.4**). The percentage of variability explained by the PCA is small (**Fig. 3**), but similar to what has been found for maize in studies focusing on different races or time periods (Arteaga et al. 2016; Rojas-Barrera et al. 2019). The admixture analysis for all races presents little genetic differentiation considering values of *K* from 2 to 4; that is, the structure is not congruent with race clustering, but rather with the geographic origin (lat/long and Pacific vs Atlantic coast) of the samples. Previous studies on the genetic structure of native maize have shown the same low levels of genetic differentiation (Pressoir & Berthaud, 2004), confirming that genetic and phenotypic variation among races is unlikely correlated (Caldu-Primo et al. 2017). This lack of genetic differentiation is attributed to the high degree of gene flow due to maize open pollination and the exchange of seeds between campesinos (Dyer & Taylor, 2008; Orozco-Ramírez et al. 2016), and also to the fact that relatively few loci are involved in the morphological traits under artificial selection (Caldú-Primo et al. 2017). Under this scenario, it is expected for gene flow to decrease population differentiation across the entire genome, with natural and artificial selection leading to local adaptations. For example, genetic diversity associated with flowering time in maize landraces varies with latitude and altitude (Romero Navarro et al. 2017). The genetic drift and natural selection processes leading to the observed patterns are independent of the race morphological characteristics (Caldu-Primo et al. 2017), which are under strong artificial selection (e.g. in this case the thin and soft cob of *Olotillo*). Our population structure analyses support this because genetic diversity within *Olotillo* and other races is influenced by the geographic origin of the samples more than by race identity (Sanchez & Goodman, 1992; Vigouroux et al. 2008). Thus, our results support what has been proposed within the maize community: maize races should be considered as morphological clusters of a set of different populations with different genetic variability at the regional level (Ortega Paczka, 2024; Ron et al. 2006; Sánchez et al. 2000).

When considering the State and scale variables the results highlight the importance of considering both the geographic scale and the local, regional, and national context when studying the genetic structure within a maize race. Further research could test if variations in genetic composition at State and geographic levels reflect local adaptation.

### Genetic diversity within the Olotillo race and signals of population contraction

As previously mentioned, earlier studies often used only a few samples per racial group, focusing on a narrow region of each race distribution; this has occurred both in genetics (e.g. Arteaga et al. 2016; Pineda-Hidalgo et al. 2013) and agronomic studies (Torres-Morales et al. 2022). Here, *Olotlillo* was extensively sampled. The genetic diversity values can not be compared to the other sampled races (*Dzit-bacal*, *Mix,* and *Tuxpeño*) due to the differences in sampling size (we include them as part of the population structure analyses). Similarly, it is not statistically fair to compare our results with those of Arteaga et al. (2016) because the sampling number and the molecular markers are different (Kumar et al. 2022).

In this study the Tajima’s D estimate indicates population contraction within *Olotillo* populations. This means that the *Olotillo* population seems to have declined over time, which could result in a smaller, more genetically uniform population with significant consequences for genetic diversity and survival. This has been mentioned by previous ethno-rural studies (Bellon M. 1996). A population decline was expected due to fewer farmers producing this race, and mainly for self-consumption (use few mother plants for seed). Additionally, those farmers who keep it only perform strong artificial selection to get the desired *Olotillo* traits and to avoid a “tuxpenized” or “hybrid-like” phenotype. This is also congruent with Bellon & Brush (1994) and with our own field observations of *Olotillo* being grown in smaller areas compared to hybrids, and even to the*Tuxpeño* race, along with intensive selection due to the search for traits appreciated in *Olotillo*. Importantly, *Olotillo* is grown close to (often much larger) fields of *Tuxpeño*, so it is possible that the number of cobs used to produce seed is further decreased in order to retain the desired phenotypic characteristics of the race. As a result, few plants are used as seed, as has been seen in several other landraces (McLean-Rodríguez et al. 2021). Thus, the inbreeding coefficient for *Olotillo* (*F*_IS_ = 0.74547 sd ± 0.06) is consistent with a contraction process, given by farmer practices (i.e. small plots, strong artificial selection) under the traditional seed system (Bellon 1991) and by changes to how much the race is being cultivated and their seeds exchanged (Bellon & Brush 1994).

Due to the genome sampling nature of GBS, nucleotide diversity may be underestimated when computed using VCFtools. Nonetheless, the heterozygosity and inbreeding indices support a pattern of inbreeding and population contraction. Specifically, high *F_IS_* values on all scales and the marked deficit of observed relative to expected heterozygosity (*Hₒ* < H*ₑ*). Similar patterns have been reported in other studies of maize genetic diversity (Badu-Apraku et al. 2021). This heterozygote deficiency can arise from several evolutionary and demographic processes. Positive *F*_IS_ values may result from non-random mating (e.g., inbreeding), directional selection against heterozygotes, population substructure (Wahlund effect), or genetic drift (De Meeûs 2018; O’Reilly et al. 2024), which again in *Olotillo* seem to be related to management practices involving relatively small fields and strong selection for the desired traits.

Additionally, population genetics statistics (*H*_e_EX, *H*_O_*EX* and *π*) also support the evidence of population decline in *Olotillo*. Notably, no significant differences were detected for these parameters across the local, regional, and national scales (**Figs. S11-S13**). This could mean that farmers undertake similar seed management across different regions from Mexico, as has been previously described by other studies (Dyer et al. 2014). This would imply that the same evolutionary processes are maintained at different scales, but does not mean that *Olotillo* populations from different parts of Mexico are the same at the genetic level, given that genetic structure was found (e.g. differences between Chiapas and Guerrero), rather, that the processes that maintain genetic variation are similar across regions of Mexico.

It is important to note that although statistically significant differences were found for *F_IS_*(**Fig. S9**) and *H_e_*OB (**Fig. S10**) at the regional and national scales, the effect sizes are modest. Despite this modest effect, greater variability in these parameters was observed, particularly at the local scale. For conservation planning, a practical starting point would be to promote seed exchange within and between communities at the local and national levels, taking into account the genetic variation present at these scales and the effect of local adaptation. The choice of encouraging exchange within and between communities will depend on how much homogenization and loss of genetic diversity are desired, with a view to obtaining local adaptation or increasing heterozygosity. Such actions could potentially reduce inbreeding and increase genetic diversity within just one or two generations.

Genomic evidence of population contraction and inbreeding in *Olotillo* supports earlier agronomic and rural observations that, although maize landraces remain widely cultivated, their local populations are shrinking. Although millions of *campesino* farmers grow maize landraces as part of both subsistence farming and responding to local and regional demand for maize (Bellon et al. 2018; 2021), there number of local varieties and the area in which they are grown has reduced over the past decades (Bellon et al. 2021; Brush & Perales, 2007; Dyer et al. 2014; McLean-Rodríguez et al. 2019), and that is being reflected at the genetic level. Several factors are driving the reduction in landrace cultivation, involving intertwined social, economical, historical, public policy and agronomical factors, like the reduced economic value of maize, farming small wages and competition from external products (Bellon et al. 2018; 2021; Monroy-Sais et al. 2024). Along with the reduction of area planted with landraces, there has been an increase of area planted with hybrid varieties, which are characterized by higher cob and grain yields whose primary destination will be commercial (McLean-Rodríguez et al. 2021), a displacement that was observed in the field. This does not mean that traditional landraces are abandoned, in fact many farmers may grow both hybrids and landraces but for different reasons and values (Monroy-Sais et al. 2024). It should be noted that for the state of Chiapas, Mexico, commercial tortillas are produced using hybrids from the state of Sinaloa, which means these hybrids do not compete with local landraces. Thus, although hybrids have not replaced landraces and likely would not do it (Monroy-Sais et al. 2024), they have contributed to a reduction in area planted with races like *Olotillo*, which in turns translates to population contraction, and farmers likely applying stronger artificial selection to keep the race characteristics, as explained above.

In summary, although *Olotillo* populations exhibit signatures of inbreeding coefficients, likely driven by recent reductions in cultivated area and limited seed exchange, substantial genetic diversity persists across the regions where the race is still maintained. This pattern highlights the importance of integrating both genetic erosion and remaining regional diversity into breeding and conservation strategies. Given the social context of *Olotillo* cultivation and consumption, conservation strategies should prioritize farming families who maintain this maize race, recognizing the importance and supporting local maize markets (Bellon et al. 2021) and breeding methods methods like evolutionary breeding (Suneson C.1956) or other participatory breeding methods (Smith et al. 2001; Vaz Patto et al. 2008). The former strategies leverage the existing diversity, prevent further genetic erosion, and have a social component. More specifically, we suggest that conservation strategies could include: (1) treating each population within the *Olotillo* race as an independent genetic unit for local breeding programs, ensuring that traits valued by farmer families are preserved; (2) designing conservation actions tailored to different geographic scales, taking into account possible local adaptation; (3) promoting and strengthening traditional farming practices and seed exchange networks to counteract genetic erosion; and (4) extending similar genetic diversity assessments to other maize races to anticipate and mitigate potential losses of genetic diversity. For *ex situ* conservation, the strengthening of local seed banks and agrobiodiversity fairs is suggested, emphasizing the importance of representing different localities and farming families within the same maize race.

## Conclusions

This study reveals fine-grain population genetics of the *Olotillo* maize race, being the first time that a large number of molecular markers have been used for a single race across all the area where it is grown. The results show that the *Olotillo* race is not a homogeneous unit but that it encompasses genetic diversity across different geographical scales, with geographic structure reflecting distances most likely used for seed exchange. Disentangling local adaptation from isolation by distance could be explored in more detail with subsequent analyses. The inbreeding values found in this study, with a large number of markers, suggest that *Olotillo* genetic pool may be undergoing size contractions despite retaining a broad distribution. Nevertheless, the fact that there are different genetic clusters within *Olotillo*, shows that even within a single maize race there are genetic differences at the population level that monitoring and conservation actions should consider, and that could be a source of adaptive potential for the future of the *Olotillo* race. Thus, populations within the *Olotillo* race should be considered as independent genetic units, tailoring conservation actions across geographic scales. Specifically, for *in situ* conservation, priority should be given to approaches such as evolutionary breeding or other participatory breeding methods and. For *ex situ* conservation, emphasis should be placed on ensuring representation of different localities and farming families within local seed banks and seed exchange fairs.

## Supporting information

All tables

## Acknowledgments

We dedicate this article to the campesinos of Mexico, especially to José Antonio Pérez López and his father, Ausencio Pérez López, guardians of the *Olotillo* in El Gavilán, Ocozocuautla. We also honor the memory of Ausencio Pérez López and Gerardo Gutiérrez Figueroa.

Thanks to Alfredo Villarruel Arroyo, Flavio Aragon Cuevas, Gerardo Gutiérrez Figueroa, Noe Orlando Gomez Montiel and Victor Antonio Vidal Martínez for support during sampling. This work was supported by CONABIO through access to its computing cluster and technical assistance by Ernesto Campos Murillo, as well as Nancy Corona and Francisca Acevedo Gasman for administrative and institucional support. We also acknowledge Mauricio R. Bellon for comments and advice, Nancy Gálvez Reyes and Marco Tulio Solano De la Cruz for laboratory assistance, INIFAP Ocozocoautla staff for fieldwork support and CIMMYT for providing part of the seed material.

This article is one of the requirements of the Programa de Doctorado en Ciencias Biológicas, Universidad Nacional Autónoma de México, for the first author to obtain a doctoral degree co-supervised by A. M-Y and A. W. D.O-G thanks the Secretaría de Ciencia, Humanidades, Tecnología e Innovación (SECIHTI) a PhD scholarship (CVU no. 545331).

## Author Contributions

D. O-G., B. C-E., H. P., A. W., and A. M-Y. conceptualized the studio. D. O-G. led the sampling with support from B. C-E. and H. P., including processing samples at INIFAP facilities. D. O-G. prepared the material, and performed the DNA extractions in lab facilities provided by D. P. and A. W. at UNAM. I. R-B. conducted the call for variants for the first plate, for a second plate D. O-G. made this. Bioinformatics analyses were made by D. O-G. and A. M-Y. with feedback from D. P. Writing was led by D. O-G. with support of A. M-Y. and A. W. All authors read and approved the final manuscript.

## Data availability

All R scripts used for this study are available in the GitHub online repository: https://github.com/Duhyadi/Paper1. SNP data and samples metadata are available at Dryad repository doiXXXXXXX (available upon acceptance). Sequence information was deposited at NCBI with project id XXXXXX(available upon acceptance).

## References

Abraham-Juárez M, Barnes A, Aragón-Raygoza A, Tyson D, Kur A, Strable J, Rellán-Álvarez R (2021) The arches and spandrels of maize domestication, adaptation, and improvement. Current Opinion in Plant Biology 64:102124. 10.1016/j.pbi.2021.102124

Alexander D, Novembre J, Lange K (2009) Fast model-based estimation of ancestry in unrelated individuals. Genome Research 19:1655–1664. https://doi,org/10,1101/gr,094052,109

Armstrong E, Mooney J, Solari K, Kim B, Barsh G, Grant V, Greenbaum G, Kaelin C, Panchenko K, Pickrell J, Rosenberg N, Ryder O, Yokoyama T, Ramakrishnan U, Petrov D, Hadly E (2024) Unraveling the genomic diversity and admixture history of captive tigers in the United States. Proceedings of the National Academy of Sciences 121(39), e2402924121. 10.1073/pnas.2402924121

Arteaga M, Moreno-Letelier A, Mastretta-Yanes A, Vázquez-Lobo A, Breña-Ochoa A, Moreno-Estrada A, Eguiarte L, Piñero D (2016) Genomic variation in recently collected maize landraces from Mexico. Genomics Data 7:38–45. https://doi,org/10,1016/j,gdata,2015,11,002

Badstue L., Bellon M, Berthaud J, Juárez X, Rosas I, Solano A, Ramírez A (2006) Examining the role of collective action in an informal seed system: a case study from the Central Valleys of Oaxaca, Mexico. Human Ecology 34:249–273. 10.1007/s10745-006-9016-2

Badu-Apraku B, Garcia-Oliveira A, Petroli C, Hearne S, Adewale S, Gedil M (2021) Genetic diversity and population structure of early and extra-early maturing maize germplasm adapted to sub-Saharan Africa. BMC Plant Biology 21:1–15. 10.1186/s12870-021-02829-6

Bellon M (1991) The ethnoecology of maize variety management: a case study from Mexico. Human Ecology 19:389–418.

Bellon M (1996) The dynamics of crop infraspecific diversity: A conceptual framework at the farmer level 1. Economic Botany 50:26–39.

Bellon M, Brush S (1994) Keepers of maize in Chiapas, Mexico. Economic Botany, 48:196–209.

Bellon M, Berthaud J, Smale M, Aguirre J, Taba S, Aragón F, Díaz J, Castro H (2003) Participatory landrace selection for on-farm conservation: An example from the Central Valleys of Oaxaca, Mexico. Genetic Resources and Crop Evolution 50:401–416.

Bellon M, Hellin J (2011) Planting hybrids, keeping landraces: agricultural modernization and tradition among small-scale maize farmers in Chiapas, Mexico. World Development 39:1434–1443. https://doi,org/10,1016/j,worlddev,2010,12,010

Bellon M, Mastretta-Yanes A, Ponce-Mendoza A, Ortiz-Santamaría D, Oliveros-Galindo O, Perales H, Acevedo F, Sarukhán J (2018) Evolutionary and food supply implications of ongoing maize domestication by Mexican campesinos. Proceedings of the Royal Society B: Biological Sciences 285:20181049. https://doi,org/10,1098/rspb,2018,1049

Bellon M, Mastretta-Yanes A, Ponce-Mendoza A, Ortiz-Santa María D, Oliveros-Galindo O, Perales H, Acevedo F, Sarukhán J (2021) Beyond subsistence: the aggregate contribution of campesinos to the supply and conservation of native maize across Mexico. Food Security 13:39–53. 10.1007/s12571-020-01134-8

Bellon M, Risopoulos J (2001) Small-scale farmers expand the benefits of improved maize germplasm: A case study from Chiapas, Mexico. World Development 29:799–811. https://doi,org/10,1016/S0305-750X(01)00013-4

Bøhn T, Aheto D, Mwangala F, Fischer K, Bones I, Simoloka C, Mbeule I, Schmidt G, Breckling B (2016) Pollen-mediated gene flow and seed exchange in small-scale Zambian maize farming, implications for biosafety assessment. Scientific Reports 6:1–12. 10.1038/srep34483

Brush S, Perales H (2007) A maize landscape: Ethnicity and agro-biodiversity in Chiapas Mexico. Agriculture, Ecosystems & Environment 121:211–221. 10.1016/j.agee.2006.12.018Get rights and content

Caballero J, Pizaña H, Cabañas A, González A, Nuñez E, Aguilar F, Ovando E (2023) Composición morfológica y rendimientos de maíces nativos sin uso de agroquímicos en Chiapas, México. Siembra 10:e3997. https://doi,org/10,29166/siembra,v10i2,3997

Caldu-Primo J, Mastretta-Yanes A, Wegier A, Piñero D (2017) Finding a needle in a haystack: distinguishing Mexican maize landraces using a small number of SNPs. Frontiers in Genetics 8:1–12. https://doi,org/10,3389/fgene,2017,00045

Callejas-Chavero A, Cornejo-Romero A, Serrato-Díaz A, Rendón-Aguilar, B (2019) Diversity and trophic structure of insects associated with grains of three maize landraces in San Agustín Loxicha, Oaxaca, Mexico. Entomological Science 22:42–47. https://doi,org/10,1111/ens,12335

CBD. (2022) Decision adopted by the Conference of the Parties to the Convention on Biological Diversity CBD/COP/DEC/15/4 Kunming-Montreal Global Biodiversity Framework. CBD/COP/DEC/15/4. https://www.cbd.int/doc/decisions/cop-15/cop-15-dec-04-en.pdf

CONABIO (2011) Proyecto Global de Maíces Nativos. Comisión Nacional para el Conocimiento y Uso de la Biodiversidad; Instituto Nacional de Investigaciones Forestales, Agrícolas y Pecuarias; Instituto Nacional de Ecología y cambio Climático. México. https://biodiversidad,gob,mx/diversidad/proyectoMaices

CONABIO (2020) Maíces https://www,biodiversidad,gob,mx/diversidad/proyectoMaices. Comisión Nacional para el Conocimiento y Uso de la Biodiversidad, Ciudad de México. México. Contenido: Cecilio Mota Cruz, Caroline Burgeff y Francisca Acevedo Gasman.

Crossa J, Taba S, Eberhart S, Bretting P, Vencovsky R (1994) Practical considerations for maintaining germplasm in maize. Theor Appl Genet 89:89–95. 10.1007/BF00226988

Cuervo-Robayo A, Ureta C, Gómez-Albores M, Meneses-Mosquera A, Téllez-Valdés O, Martínez-Meyer E (2020) One hundred years of climate change in Mexico. PLoS ONE 15:1–19. 10.1371/journal.pone.0209808

Danecek P, Auton A, Abecasis G, Albers C, Banks E, DePristo M, Handsaker R, Lunter G, Marth G, Sherry S, McVean G, Durbin R, 1000 Genomes Project Analysis Group (2011) The variant call format and VCFtools. Bioinformatics 27:2156–2158. 10.1093/bioinformatics/btr330

De Meeûs T. (2018). Revisiting F_IS_, F_ST_, Wahlund effects, and null alleles. Journal of Heredity, 109:446–456. 10.1093/jhered/esx106

Dyer G, López-Feldman A (2013) Inexplicable or simply unexplained? The management of maize seed in Mexico. PLoS One 8:e68320. 10.1371/journal.pone.0068320

Dyer G., López-Feldman A., Yúnez-Naude A, Taylor E (2014) Genetic erosion in maize’s center of origin. Proceedings of the National Academy of Sciences 111:14094–14. 10.1073/pnas.1407033111

Dyer G, Taylor E (2008) A crop population perspective on maize seed systems in Mexico. Proceedings of the National Academy of Sciences 105:470–475. 10.1073/pnas.0706321105

Goodman M, Paterniani E (1969) The races of maize: III. Choices of appropriate characters for racial classification. Economic Botany 23:265–273. 10.1007/BF02860459

Hemstrom W, Grummer J, Luikart G, Christie M (2024) Next-generation data filtering in the genomics era. Nature Reviews Genetics 25:750–767. 10.1038/s41576-024-00738-6

Hellin J, Bellon M, Hearne S (2014) Maize landraces and adaptation to climate change in Mexico. Journal of Crop Improvement 28:484–501. 10.1080/15427528.2014.921800

Herten K, Hestand M, Vermeesch J, Van Houdt J (2015) GBSX: a toolkit for experimental design and demultiplexing genotyping by sequencing experiments. BMC Bioinformatics 16:1–6. 10.1186/s12859-015-0514-3

Korunes K, Samuk K (2021) pixy: Unbiased estimation of nucleotide diversity and divergence in the presence of missing data. Molecular Ecology Resources, 21:1359–1368. 10.1111/1755-0998.13326

Kumar A, Longmei N, Kumar P, Kaushik P (2022) Molecular marker analysis of genetic diversity in maize: A review. OBM Genetics 6:1–19. 10.21926/obm.genet.2201150

Li H, Handsaker B, Wysoker A, Fennell T, Ruan J, Homer N, Marth G, Abecasis G, Durbin R, 1000 Genome Project Data Processing Subgroup (2009) The sequence alignment/map format and SAMtools. Bioinformatics 25:2078–2079. 10.1093/bioinformatics/btp352

Llamas-Guzmán L, Lazos E, Perales H, Casas A (2022) Seed Exchange Networks of Native Maize, Beans, and Squash in San Juan Ixtenco and San Luis Huamantla, Tlaxcala, Mexico. Sustainability 14:1–34. 10.3390/su14073779

López-Morales F, Vázquez-Carrillo M, Molina-Galán J, García-Zavala J, Corona-Torres T, Cruz-Izquierdo S, López-Romero G, Reyes-López D, Esquivel-Esquivel G (2017) Interacción genotipo-ambiente, estabilidad del rendimiento y calidad de grano en maíz Tuxpeño. Revista Mexicana de Ciencias Agrícolas 8:1035–1050. 10.29312/remexca.v8i5.106

Mastretta-Yanes A, Tobin D, Bellon M, von Wettberg E, Cibrián-Jaramillo A, Wegier A, Monroy-Sais A, Galvez-Reyes N, Ruiz-Arrocho J, Chen Y (2024) Human management of ongoing evolutionary processes in agroecosystems. Plants, People, Planet 6:1190–1206.

McKenna A, Hanna M, Banks E, Sivachenko A, Cibulskis K, Kernytsky A, Garimella K, Altshuler D, Gabriel S, Daly M, DePristo M (2010) The Genome Analysis Toolkit: a MapReduce framework for analyzing next-generation DNA sequencing data. Genome research, 20:1297–1303. 10.1101/gr.107524.110

McLean-Rodríguez F, Camacho-Villa T, Almekinders C, Pè M, Dell’Acqua M, Costich D (2019) The abandonment of maize landraces over the last 50 years in Morelos, Mexico: a tracing study using a multi-level perspective. Agric and Hum Values 36:651–668. 10.1007/s10460-019-09932-3

McLean-Rodríguez F, Costich D, Camacho-Villa T, Pè M, Dell’Acqua M (2021) Genetic diversity and selection signatures in maize landraces compared across 50 years of in situ and ex situ conservation. Heredity 126:913–928. 10.1038/s41437-021-00423-y

Mercer K, Perales H (2019) Structure of local adaptation across the landscape: flowering time and fitness in Mexican maize (*Zea mays* L, subsp, mays) landraces. Genetic Resour Crop Evol 66:27–45. 10.1007/s10722-018-0693-7

Mercer K, Wainwright J (2008) Gene flow from transgenic maize to landraces in Mexico: an analysis. Agriculture, ecosystems & environment 123:109–115. 10.1016/j.agee.2007.05.007

Monroy-Sais A, Tobin D, Bellon M, Astier M, Cibrián-Jaramillo A, Gálvez-Reyes N, Mastretta-Yanes A, Ruiz-Arrocho J, Wegier A, Chen Y (2024) Smallholder farmers’ diverse values in maize landrace conservation: A case study from Chiapas, Mexico. Journal of Rural Studies 110:103347. 10.1016/j.jrurstud.2024.103347

Nuss E, Tanumihardjo S (2010) Maize: a paramount staple crop in the context of global nutrition. Comprehensive reviews in food science and food safety 9: 417–436. 10.1111/j.1541-4337.2010.00117.x

Ocampo-Giraldo V, Camacho-Villa C, Costich D, Vidal Martínez V, Smale M, Jamora N (2020) Dynamic conservation of genetic resources: Rematriation of the maize landrace Jala. Food Security 12:945–958. 10.1007/s12571-020-01054-7

Orozco-Ramírez Q, Ross-Ibarra J, Santacruz-Varela, A, Brush S (2016) Maize diversity associated with social origin and environmental variation in Southern Mexico. Heredity 116:477–484. 10.1038/hdy.2016.10

Ortega Packa (2024) Cultivo de maíces nativos en México y qué hace falta para caracterizarlos. In: “Diálogos por la transformación: el maíz nativo”. Universidad Rosario Castellanos. Ciudad de México, 20 de junio.

O’Reilly G, Manlik, O, Vardeh S, Sinclair J, Cannell B, Lawler Z., Sherwin W (2024). A new method for ecologists to estimate heterozygote excess and deficit for multi-locus gene families. Ecology and Evolution, 14:e11561. 10.1002/ece3.11561

Patil I (2021) Visualizations with statistical details: The ‘ggstatsplot’ approach. Journal of Open Source Software 6: 3167. 10.21105/joss.03167

Palacios-Pola G, Perales H, Estrada E, Figueroa-Cárdenas J, D, D (2022) Nixtamal techniques for different maize races prepared as tortillas and tostadas by women of Chiapas, Mexico. Journal of Ethnic Foods 9:1–10. 10.1186/s42779-022-00116-9

Prasanna B (2012) Diversity in global maize germplasm: characterization and utilization. Journal of biosciences 37:843–855. 10.1007/s12038-012-9227-1

Perales H, Golicher D (2014) Mapping the diversity of maize races in Mexico. PloS One 9:1–20. 10.1371/journal.pone.0114657

Perales H, Hernández (2005) Diversidad del maíz en Chiapas. In: González-Espinosa, M.; Ramírez-Marcial N. and Ruiz-Montoya L. (Coord.). Diversidad biológica en Chiapas. Plaza y Valdés, SA de CV. México, DF. 419–440 pp.

Pineda-Hidalgo K., Méndez-Marroquín K, Vega E, Chávez-Ontiveros J, Sánchez-Peña P, Garzón-Tiznado J, Vega-García M, López-Valenzuela J (2013) Microsatellite-based genetic diversity among accessions of maize landraces from Sinaloa in México. Hereditas, 150:53–59. 10.1111/j.1601-5223.2013.00019.x

Price A, Patterson N, Plenge R, Weinblatt M, Shadick N, Reich D (2006) Principal components analysis corrects for stratification in genome-wide association studies. Nat Genet 38:904–909. 10.1038/ng1847

Pressoir G, Berthaud J (2004) Population structure and strong divergent selection shape phenotypic diversification in maize landraces. Heredity 92(2):95–101. 10.1038/sj.hdy.6800388

Purugganan M (2022) What is domestication?. Trends in ecology & evolution 37:663–671. : 10.1016/j.tree.2022.04.006

R Core Team (2024) *R 4.4.1*: A Language and Environment for Statistical Computing. R Foundation for Statistical Computing, Vienna, Austria. https://www.R-project.org/

R Core Team (2025) *R 4.5.0:* A Language and Environment for Statistical Computing. R Foundation for Statistical Computing, Vienna, Austria. https://www.R-project.org/

Rice E (2007) Conservation in a changing world: in situ conservation of the giant maize of Jala. Genet Resour Crop Evol 54:701–713. 10.1007/s10722-006-0023-3

Rocandio-Rodríguez M, Santacruz-Varela A, Córdova-Téllez L, López-Sánchez H, Castillo-González F, Lobato-Ortiz R, García-Zavala J (2014) Detection of genetic diversity of seven maize races from the high central valleys of Mexico using microsatellites. Maydica 59:144–151.

Rojas-Barrera I, Wegier A, Sánchez J, Owens G, Rieseberg L, Piñero D (2019) Contemporary evolution of maize landraces and their wild relatives influenced by gene flow with modern maize varieties. Proceedings of the National Academy of Sciences, 116:21302–21311. 10.1073/pnas.1817664116

Romero Navarro J, Willcox M, Burgueño J, Romay C, Swarts K, Trachsel S, Preciado E, Terron A, Vallejo Delgado H, Vidal V, Ortega A, Espinoza Banda A, Gómez Montiel O, Ortiz-Monasterio I, San Vicente F, Guadarrama Espinoza A, Atlin G, Wenzl P, Hearne S, Buckler E (2017) A study of allelic diversity underlying flowering-time adaptation in maize landraces. Nat Genet 49:476–480. 10.1038/ng.3784

Ron J, Sánchez J, Jiménez A, Carrera, J, Martín J, Morales M, de la Cruz L, Hurtado S, Munguía S, Rodríguez J (2006) Maíces nativos del occidente de México I. Colectas 2004. Scientia-CUCBA 8:1–139.

Ruiz J, Durán N, Sanchez J, Ron P, Gonzalez D, Holland J, Medina G (2008) Climatic adaptation and ecological descriptors of 42 Mexican maize races. Crop Science 48:1502–1512. 10.2135/cropsci2007.09.0518

Sanchez J, Goodman M (1992) Relationships among the Mexican races of maize. Econ Bot 46:72–85. 10.1007/BF02985256

Sánchez J, Goodman M, Rawlings O (1993) Appropriate characters for racial classification in maize. Econ Bot 47:44–59.

Sanchez J, Goodman M, Stuber C (2000) Isozymatic and morphological diversity in the races of maize of Mexico. Econ Bot 54:43–59.

Sedlazeck F, Rescheneder P, von Haeseler A (2013) NextGenMap: fast and accurate read mapping in highly polymorphic genomes. Bioinformatics 29:2790–2791. 10.1093/bioinformatics/btt468

Sierra-Macías M, Andrés-Meza P, Rodríguez-Montalvo F, Espinosa Calderon A (2016) Introgresión Genética de genotipos mejorados en maíces nativos de las razas Tuxpeño y Olotillo con calidad de hoja del totomoxtle. Revista de Aplicación Científica y Técnica 2:45–52.

Smith M, Castillo F, Gómez F (2001) Participatory plant breeding with maize in Mexico and Honduras. Euphytica, 122, 551–563. 10.1023/A:1017510529440

Staller J (2009) Maize cobs and cultures: history of Zea mays L. Springer Science & Business Media.

Suneson C (1956) An evolutionary plant breeding method 1. Agronomy Journal 48:188–191.

Torres-Morales B, Rocandio-Rodríguez M, Santacruz-Varela M, Córdova-Téllez L, Coutiño-Estrada B, López-Sánchez Higinio (2022) Morphological and agronomic diversity corn races from the state of Chiapas. Revista Mexicana de Ciencias Agrícolas 13:687–699. 10.29312/remexca.v13i4.2956

Van Heerwaarden J, Hellin J, Visser R, Van Eeuwijk (2009) Estimating maize genetic erosion in modernized smallholder agriculture. Theoretical and Applied Genetics 119:875–888. 10.1007/s00122-009-1096-0

Vaz Patto M, Moreira P, Almeida N, Satovic Z, Pego S (2008) Genetic diversity evolution through participatory maize breeding in Portugal. Euphytica 161:283–291. 10.1007/s10681-007-9481-8

Vigouroux Y., Glaubitz J., Matsuoka Y., Goodman M., Sánchez J., Doebley J (2008) Population structure and genetic diversity of New World maize races assessed by DNA microsatellites. American Journal of Botany, 95:1240–1253. 10.3732/ajb.0800097

Wang N, Yuan Y, Wang H, Yu D, Liu Y, Zhang A, Gowda M, Nair S, Hao H, Lu Y, San Vicente F, Prassana B, Li X, Zhang, X (2020) Applications of genotyping-by-sequencing (GBS) in maize genetics and breeding. Sci Rep 10:1–12. 10.1038/s41598-020-73321-8

Wellhausen E, Roberts L, Hernández E (1951) Razas de maíz en México, su origen, características y distribución. Folleto técnico núm, 5. Oficina de Estudios Especiales (OEE), Secretaría de Agricultura y Ganadería (SAG), México. DF. 237 p.

Wertz S (2005) Maize: the native North American’s legacy of cultural diversity and biodiversity. Journal of Agricultural and Environmental Ethics, 18:131–156. DOI 10.1007/s10806-005-0635-1

Zimmerman S, Aldridge C, Oyler-McCance S (2020) An empirical comparison of population genetic analyses using microsatellite and SNP data for a species of conservation concern. BMC Genomics 21:1–16. 10.1186/s12864-020-06783-9

Zheng X, Levine D, Shen J, Gogarten S, Laurie C, Weir B (2012) A high-performance computing toolset for relatedness and principal component analysis of SNP data. Bioinformatics 28:3326–3328. 10.1093/bioinformatics/bts606

