## Supplementary material for "Diversity and genetic structure within a Mexican maize race reveal consistent biocultural processes across geographic scales": TableS1

| ID | collector_name | source | accesion_bank | collection_year | country | state | municipality | locality | latitude | longitude | altitude | species | race | scale | plant_material |
|---|---|---|---|---|---|---|---|---|---|---|---|---|---|---|---|
| CHIS_R02 | Duhyadi Oliva Garcia | field | | 2018 | Mexico | Chiapas | Villaflores | Ricardo Flores Magón | 16.299 | -93.592 | 731 | *Zea mays* ssp. *mays* | *Olotillo* | regional | 100_ears |
| CHIS_L04 | Duhyadi Oliva Garcia | field | | 2018 | Mexico | Chiapas | Ocozocuatla | El Gavilán | 16.749 | -93.458 | 758 | *Zea mays* ssp. *mays* | *Olotillo* | local | 100_ears |
| CHIS_R05 | Bulmaro Coutiño Estrada | field | | 2018 | Mexico | Chiapas | Cintalapa | Corazón del Valle | 16.416 | -93.983 | 810 | *Zea mays* ssp. *mays* | *Olotillo* | regional | 100_ears |
| CHIS_R06 | Duhyadi Oliva Garcia | field | | 2018 | Mexico | Chiapas | Suchiapa | El Recreo "Plan de Mulumi" | 16.634 | -93.121 | 455 | *Zea mays* ssp. *mays* | *Olotillo* | regional | 130_ears |
| CHIS_L08 | Bulmaro Coutiño Estrada | field | | 2018 | Mexico | Chiapas | Ocozocuatla | San Isidro, Ciénega Silviana_ | 16.67667 | -93.41167 | 786 | *Zea mays* ssp. *mays* | *Olotillo* | local | 100_ears |
| CHIS_R10 | Duhyadi Oliva Garcia | field | | 2018 | Mexico | Chiapas | Cintalapa | Villamorelos | 16.478 | -93.928 | 711 | *Zea mays* ssp. *mays* | *Olotillo* | regional | 100_ears |
| CHIS_R11 | Duhyadi Oliva Garcia | field | | 2018 | Mexico | Chiapas | El Parral | El Parral | 16.364 | -93.001 | 650 | *Zea mays* ssp. *mays* | *Olotillo* | regional | 100_ears |
| GRO_N13 | Noel Orlando Gomez Montiel | field | | 2018 | Mexico | Guerrero | San Luis Acatl | Yoloxochitl | 16.8115278 | -98.68422222 | 610 | *Zea mays* ssp. *mays* | *Olotillo* | national | 20_ears |
| GRO_N14 | Noel Orlando Gomez Montiel | field | | 2018 | Mexico | Guerrero | San Luis Acatl | El Carmen | 16.8155 | -98.76111111 | 287 | *Zea mays* ssp. *mays* | *Olotillo* | national | 20_ears |
| GRO_N15 | Noel Orlando Gomez Montiel | field | | 2018 | Mexico | Guerrero | San Luis Acatl | Cuanacaxtitlan | 16.7951944 | -98.63719444 | 438 | *Zea mays* ssp. *mays* | *Olotillo* | national | 20_ears |
| GRO_N16 | Noel Orlando Gomez Montiel | field | | 2018 | Mexico | Guerrero | San Luis Acatl | Yoloxochitl | 16.8158056 | -98.68438889 | 615 | *Zea mays* ssp. *mays* | *Olotillo* | national | 20_ears |
| GRO_N18 | Noel Orlando Gomez Montiel | field | | 2018 | Mexico | Guerrero | Coyuca de Be | Atoyaquillo | 17.1046944 | -100.0449722 | 175 | Zea mays ssp. mays | Olotillo | national | 10_ears |
| GRO_N19 | Noel Orlando Gomez Montiel | field | | 2018 | Mexico | Guerrero | San Luis Acatl | Yoloxochitl | 16.8155 | -90.68991667 | 615 | *Zea mays* ssp. *mays* | *Olotillo* | national | 20_ears |
| GRO_N20 | Noel Orlando Gomez Montiel | field | | 2018 | Mexico | Guerrero | Coyuca de Be | Pueblo Viejo | 17.0998889 | -99.99258333 | 520 | *Zea mays* ssp. *mays* | *Olotillo* | national | 10_ears |
| GRO_N22 | Noel Orlando Gomez Montiel | field | | 2018 | Mexico | Guerrero | San Luis Acatl | Yoloxochitl | 16.8115278 | -98.68405556 | 641 | *Zea mays* ssp. *mays* | *Olotillo* | national | 20_ears |
| GRO_N23 | Noel Orlando Gomez Montiel | field | | 2018 | Mexico | Guerrero | Coyuca de Be | La Lima | 16.9850833 | -99.86388889 | 324 | *Zea mays* ssp. *mays* | *Olotillo* | national | 10_ears |
| GRO_N24 | Noel Orlando Gomez Montiel | field | | 2018 | Mexico | Guerrero | Coyuca de Be | Pueblo Viejo | 17.0221111 | -99.98516667 | 520 | *Zea mays* ssp. *mays* | *Olotillo* | national | 10_ears |
| GRO_N26 | Noel Orlando Gomez Montiel | field | | 2018 | Mexico | Guerrero | Coyuca de Be | Paso Real | 17.08425 | -100.0758333 | 136 | *Zea mays* ssp. *mays* | *Olotillo* | national | 10_ears |
| NAY_N27 | Víctor Antonio Vidal Martínez | field | | 2019 | Mexico | Nayarit | Ixtlán del Río | Cacalutan | 21.1166833 | -104.2615556 | 902 | *Zea mays* ssp. *mays* | *Olotillo* | national | 50_ears_&_2_kg_seeds |
| NAY_N29 | Víctor Antonio Vidal Martínez | field | | 2019 | Mexico | Nayarit | Ixtlán del Río | Los Sauces, Cacalutan | 21.1324167 | -104.2629167 | 798 | *Zea mays* ssp. *mays* | *Olotillo* | national | 66_ears |
| NAY_N30 | Víctor Antonio Vidal Martínez | field | | 2019 | Mexico | Nayarit | La Yesca | Puente de Camotlán | 21.7060833 | -104.0749444 | 1136 | *Zea mays* ssp. *mays* | *Olotillo* | national | 77_ears |
| OAX_N31 | Flavio Aragon Cuevas | field | | 2018 | Mexico | Oaxaca | Villa de Tutute | Duva-Yoo | 16.162556 | -97. 526056 | 512 | *Zea mays* ssp. *mays* | *Olotillo* | national | 100_ears |
| OAX_N32 | Flavio Aragon Cuevas | field | | 2018 | Mexico | Oaxaca | San Pedro Po | San Roque | 15.78819195 | -96.46164785 | 182 | *Zea mays* ssp. *mays* | *Olotillo* | national | 100_ears |
| CHIS_L33 | Bulmaro Coutiño Estrada | field | | 2018 | Mexico | Chiapas | Ocozocuatla | Rivera "El Gavilán" | 16.75153 | -93.45709 | 757 | *Zea mays* ssp. *mays* | *Olotillo* | local | 100_ears |
| CHIS_L34 | Bulmaro Coutiño Estrada | field | | 2018 | Mexico | Chiapas | Ocozocuatla | Ejido San Rafael | 16.666931 | -93.418475 | 785 | *Zea mays* ssp. *mays* | *Olotillo* | local | 10_ears_&_1_kg_seeds |
| CHIS_L35 | Duhyadi Oliva Garcia | field | | 2018 | Mexico | Chiapas | Ocozocuatla | El Gavilán | 16.74977 | -93.46063 | 756 | *Zea mays* ssp. *mays* | *Olotillo* | local | 100_ears |
| CHIS_L36 | Duhyadi Oliva Garcia | field | | 2019 | Mexico | Chiapas | Jiquipilas | Benito Juárez | 16.782556 | -93.608043 | 751 | *Zea mays* ssp. *mays* | *Olotillo* | local | 7_kg_seeds |
| CHIS_L37 | Duhyadi Oliva Garcia | field | | 2019 | Mexico | Chiapas | Ocozocuatla | San Jorge km 18 | 16.7129861 | -93.52033333 | 784 | *Zea mays* ssp. *mays* | *Olotillo* | local | 7_kg_seeds |
| CHIS_L38 | Duhyadi Oliva Garcia | field | | 2019 | Mexico | Chiapas | Ocozocuatla | El Aguacero | 16.780436 | -93.492936 | 775 | *Zea mays* ssp. *mays* | *Mix* | local | 1_kg_seeds |
| CHIS_R40 | Duhyadi Oliva Garcia | field | | 2018 | Mexico | Chiapas | Chiapa de Co | Narciso Mendoza | 16.57778189 | -92.98854284 | 435 | *Zea mays* ssp. *mays* | *Olotillo* | regional | 7_kg_seeds |
| CHIS_L41 | Bulmaro Coutiño Estrada | field | | 2018 | Mexico | Chiapas | Ocozocuatla | Rancho El Pitutal | 16.70745 | -93.32761 | 932 | *Zea mays* ssp. *mays* | *Olotillo* | local | 100_ears |
| CHIS_L42 | Bulmaro Coutiño Estrada | field | | 2018 | Mexico | Chiapas | Ocozocuatla | Rancho San Isidro | 16.8562 | -93.40551 | 972 | *Zea mays* ssp. *mays* | *Olotillo* | local | 100_ears |
| CHIS_L43 | Bulmaro Coutiño Estrada | field | | 2018 | Mexico | Chiapas | Ocozocuatla | Región_Mazotho_Ejido_Ocoz | 16.757344 | -93.4547 | 754 | *Zea mays* ssp. *mays* | *Olotillo* | local | 100_ears |
| CHIS_L44 | Bulmaro Coutiño Estrada | field | | 2018 | Mexico | Chiapas | Ocozocuatla | San_Jorge km 18 | 16.71199 | -93.51664 | 810 | *Zea mays* ssp. *mays* | *Olotillo* | local | 100_ears |
| CHIS_L45 | Bulmaro Coutiño Estrada | field | | 2018 | Mexico | Chiapas | Ocozocuatla | La_Borcelana_Ejido_Ocozocu | 16.78669 | -93.43072 | 841 | *Zea mays* ssp. *mays* | *Olotillo* | local | 100_ears |
| NAY_N47 | Víctor Antonio Vidal Martínez | field | | 2019 | Mexico | Nayarit | La Yesca | Huajimic | 21.6436944 | -104.3425556 | 1211 | *Zea mays* ssp. *mays* | *Olotillo* | national | 100_ears |
| VER_E48 | Victor Hugo Chavez | bank | CIMMYTMA29569 | 2007 | Mexico | Veracruz | Tantoyuca | El Mezquite | 21.3833 | -98.25 | 126 | *Zea mays* ssp. *mays* | *Olotillo* | national | 100_ears |
| SLP_E49 | Victor Hugo Chavez | bank | CIMMYTMA29498 | 2007 | Mexico | San Luis Poto | Xilitla | Apetzco | 21.393081 | -99.010927 | 872 | *Zea mays* ssp. *mays* | *Olotillo* | national | 100_ears |
| HGO_E50 | Victor Hugo Chavez | bank | CIMMYTMA29343 | 2007 | Mexico | Hidalgo | Jaltocán | La Ilusión | 21.141304 | -98.54596 | 248 | *Zea mays* ssp. *mays* | *Olotillo* | national | 100_ears |
| SLP_E51 | Victor Hugo Chavez | bank | CIMMYTMA29464 | 2007 | Mexico | San Luis Poto | Tampacan | Chupadero | 21.4 | -98.73 | 148 | *Zea mays* ssp. *mays* | *Olotillo* | national | 100_ears |
| CHIS_E52 | Hugo Perales Rivera | field | | | Mexico | Chiapas | Comitán de D | Santa Elena Las Agujas | 16.36120704 | -92.17921108 | 2089 | *Zea mays* ssp. *mays* | *Olotillo* | regional | 100_ears |
| ROO_E53 | Hugo Perales Rivera | field | | 2016 | Mexico | Quintana Roo | Felipe Carrillo | Chumpón | 20.00222 | -87.81222 | 7 | *Zea mays* ssp. *mays* | *Dzit-bacal* | national | 100_ears |
| CAM_E54 | Hugo Perales Rivera | field | | 2015 | Mexico | Campeche | Puerto Campeche | | | | | *Zea mays* ssp. *mays* | *Dzit-bacal* | national | 100_ears |
| CAM_E55 | Hugo Perales Rivera | field | | 2016 | Mexico | Campeche | Champoton | Pustunich | 19.14444 | -90.47861 | 55 | *Zea mays* ssp. *mays* | *Dzit-bacal* | national | 100_ears |
| VER_E56 | Víctor Hugo Chavez | bank | CIMMYTMA29586 | 2007 | Mexico | Veracruz | Benito Juárez | Azoquitipa | 20.77 | -98.18 | 458 | *Zea mays* ssp. *mays* | *Olotillo* | national | 100_ears |
| VER_E57 | Víctor Hugo Chavez | bank | CIMMYTMA29535 | 2007 | Mexico | Veracruz | Ixcatepec | Ejido Ixcatepec | 21.2 | -97.98 | 299 | *Zea mays* ssp. *mays* | *Olotillo* | national | 100_ears |
| CAM_E58 | Hugo Perales Rivera | field | | 2016 | Mexico | Campeche | Escárcega | Ejido La Victoria | 18.4931 | -90.9221 | 100 | *Zea mays ssp. mays* | *Tuxpeño* | national | 100_ears |
| CHIS_E59 | Hugo Perales Rivera | field | | | Mexico | Chiapas | Comitán de D | Santa Elena Las Agujas | 16.36120704 | -92.17921108 | 2089 | *Zea mays ssp. mays* | *Olotillo* | regional | 100_ears |
| CAM_E60 | Hugo Perales Rivera | field | | 2016 | Mexico | Campeche | Champoton | Pixoyal | 18.937363 | -90.599887 | 100 | *Zea mays ssp. mays* | *Tuxpeño* | regional | 100_ears |

| variable | description |
|---|---|
| ID | unique identifier for each accesion collected. |
| collector_name | full name of the person who collected the sample. |
| source | indicates whether the sample was collected directly from the field or another source, like a gene bank. |
| accesion_bank | code or label used to track the sample in a germplasm bank (CIMMYT). |
| collection_year | year in which the accesion was collected. |
| country | country in which the collection was carried out. |
| state | name of the Mexican state where the sample was collected. |
| municipality | name of the municipality within the state, administrative division. |
| locality | specific locality, name, or site where the accesion was collected. |
| latitude | geographic latitude coordinate of the collection site in decimal degrees. When the sampling place was a farmer's house, the coordiantes were generalized to obsecure the exact location for privacy reasons. |
| longitude | geographic longitude coordinate of the collection site in decimal degrees. When the sampling place was a farmer's house, the coordiantes were generalized to obsecure the exact location for privacy reasons. |
| altitude | elevation in meters above sea level at the collection site. |
| species | scientific name of the maize accesion collected: *Zea mays* ssp. *mays.* |
| race | traditional racial classification of the maize, based on farmer knowledge or phenotype. |
| scale | spatial level of sampling: local, regional, or national. |
| observations_mixed | refers to the varieties that make up the mixture. |
| plant_material | quantity of plant material collected per sample, expressed either as the number of ears (e.g., 100, 50, or 20 ears) or as the weight of seeds in kilograms, depending on the collection method used. |
