## Supplementary material for "Diversity and genetic structure within a Mexican maize race reveal consistent biocultural processes across geographic scales": All tables

**Table 1** Number of samples and accessions by State

| Sampling state | Accessions | Number of samples | Olotillo samples (*n*) | Tuxpeño samples (*n*) | Dzit-bacal samples (*n*) | Mix samples (*n*) |
|---|---|---|---|---|---|---|
| Chiapas-Local | 13 | 53 | 50 | 0 | 0 | 3 |
| Chiapas-Regional | 8 | 28 | 28 | 0 | 0 | 0 |
| Campeche | 4 | 8 | 0 | 4 | 4 | 0 |
| Guerrero | 11 | 43 | 43 | 0 | 0 | 0 |
| Hidalgo | 1 | 2 | 2 | 0 | 0 | 0 |
| Nayarit | 4 | 16 | 16 | 0 | 0 | 0 |
| Oaxaca | 2 | 9 | 9 | 0 | 0 | 0 |
| Quintana Roo | 1 | 2 | 0 | 0 | 2 | 0 |
| San Luis Potosí | 2 | 4 | 4 | 0 | 0 | 0 |
| Veracruz | 3 | 6 | 6 | 0 | 0 | 0 |
| **Total** | **49** | **171** | **158** | **4** | **6** | **3** |

[a] accessions refer to each of the collections made. Samples refer to each of the seeds germinated by accession

**Table 2** Population genetics statistics by race

| Statistic[a] | Dzit-bacal (n = 6) | Mix (n = 3) | Olotillo (n = 158) | Tuxpeño (n = 4) | All races mean (n = 171) |
|---|---|---|---|---|---|
| Mean $F_{IS}$ | 0.724 ± 0.019 | 0.771 ± 0.011 | 0.745 ± 0.067 | 0.769 ± 0.016 | 0.746 ± 0.066 |
| Median $F_{IS}$ | 0.729 | 0.773 | 0.749 | 0.771 | |
| (min, max) | (0.694, 0.740) | (0.759, 0.781) | (0.458, 1.0) | (0.750, 0.785) | |
| Mean $H_e$OB | 0.082 ± 0.006 | 0.068 ± 0.003 | 0.075 ± 0.020 | 0.068 ± 0.005 | 0.075 ± 0.019 |
| Median $H_e$OB | 0.080 | 0.067 | 0.074 | 0.068 | |
| (min, max) | (0.077, 0.090) | (0.065, 0.071) | (0, 0.161) | (0.064, 0.074) | |
| Mean $H_e$EX | 0.296 ± 0.000 | 0.295 ± 0.00 | 0.295 ± 0.002 | 0.295 ± 0.000 | 0.295 ± 0.002 |
| Median $H_e$EX | 0.296 | 0.295 | 0.296 | 0.295 | |
| (min, max) | (0.296, 0.296) | (0.295, 0.296) | (0.276, 0.296) | (0.295, 0.295) | |
| Mean $H_o$EX | 0.704 ± 0.000 | 0.705 ± 0.000 | 0.705 ± 0.002 | 0.705 ± 0.00 | 0.705 ± 0.002 |
| Median $H_o$EX | 0.704 | 0.705 | 0.704 | 0.705 | |
| (min, max) | (0.704, 0.704) | (0.704, 0.705) | (0.704, 0.724) | (0.705, 0.705) | |
| Mean $\pi$ | 1.140 e-04 ± 1.025 e-04 | 1.222 e-04 ± 1.034 e-04 | 1.013 e-04 ± 9.574 e-05 | 1.133 e-04 ± 9.920 e-05 | 1.01 e-04 ± 9.6 e-05 |
| Median $\pi$ | 8.333 e-05 | 1.000 e-04 | 7.183 e-05 | 8.571 e-05 | |
| (min, max) | (4.546 e-06, 1.159 e-03) | (6.667 e-06, 1.107 e-03) | (5.229 e-06, 9.693 e-04) | (3.572 e-06, 1.157 e-03) | |
| Mean $D$ | 0.572 ± 8.502 e-01 | 0.733 ± 7.411 e-01 | 1.357 ± 9.628 e-01 | 0.644 ± 7. 710 e-01 | 1.37 ± 9.67 e-01 |
| Median $D$ | 0.643 | 0.851 | 1.378 | 0.678 | |
| (min, max) | (-1.944, 2.539) | (-1.461, 2.238) | (-1.002, 4.962) | (-1.770, 2.359) | |
| Mean $N_s$ | 85,006 ± 428 | 83,092 ± 842 | 82,144 ± 12,236 | 83,076 ± 438 | 82,269 ± 11, 820 |
| Median $N_s$ | 85,041 | 82,714 | 84,130 | 83,131 | |
| (min, max) | (84346, 85532) | (82505, 84058) | (2186, 039) | (82492, 83550) | |

[a] $F_{IS}$ Inbreeding coefficient, $H_e$OB Observed heterozygosity, $H_e$Ex Expected heterozygosity, $H_o$Ex Expected homozygosity, $\pi$ Nucleotide diversity, $D$ Tajima's D value, and $N_s$ Number of sites

**Table 3** Population genetics statistics by scale for *Olotillo*

| Statistic[a] | Local (n =50) | Regional (n =28) | National (n =80) |
|---|---|---|---|
| Mean $F_{IS}$<br>Median $F_{IS}$<br>(min, max) | 0.715 ± 0.088<br>0.740<br>(0.458, 0.995) | 0.759 ± 0.050<br>0.759<br>(0.691, 0.880) | 0.750 ± 0.045<br>0.743<br>(0.682, 1) |
| Mean $H_e$OB<br>Median $H_e$OB<br>(min, max) | 0.084 ± 0.026<br>0.077<br>(0.001, 0.161) | 0.071 ± 0.015<br>0.071<br>(0.035, 0.091) | 0.074 ± 0.013<br>0.076<br>(0, 0.094) |
| Mean $H_e$EX<br>Median $H_e$EX<br>(min, max) | 0.296 ± 0.000<br>0.296<br>(0.295, 0.296) | 0.296 ± 0.000<br>0.296<br>(0.295, 0.296) | 0.295 ± 0.002<br>0.296<br>(0.276, 0.296) |
| Mean $H_oEX$<br>Median $H_oEX$<br>(min, max) | 0.704 ± 0.000<br>0.704<br>(0.704, 0.705) | 0.704 ± 0.000<br>0.704<br>(0.704, 0.705) | 0.705 ± 0.002<br>0.704<br>(0.704, 0.724) |
| Mean $\pi$<br>Median $\pi$<br>(min, max) | 1.015 e-04 ± 9.699 e-05<br>7.213 e-05<br>(1.420 e-06, 1.103 e-03) | 1.036 e-04 ± 9.903 e-05<br>7.349 e-05<br>(2.748 e-06, 1.057 e-03) | 1.036 e-04 ± 9.831 e-05<br>7.353 e-05<br>(9.159 e-07, 1.034 e-03) |
| Mean *D*<br>Median *D*<br>(min, max) | 0.966 ± 9.466 e-05<br>1.001<br>(-1.688, 4.214) | 0.705 ± 9.314 e-01<br>0.736<br>(-2.174, 3.601) | 1.152 ± 9.489 e-01<br>1.182<br>(-1.734, 4.614) |
| Mean $N_s$<br>Median $N_s$<br>(min, max) | 82,529 ± 11,165<br>84, 118<br>(5492, 85805) | 83,405 ± 2,637<br>83, 808<br>(72190, 85867) | 83,293 ± 9,484<br>84, 450<br>(21, 86039) |

[a] $F_{IS}$ Inbreeding coefficient, $H_e$OB Observed heterozygosity, $H_e$Ex Expected heterozygosity, $H_o$Ex Expected homozygosity, $\pi$ Nucleotide diversity, *D* Tajima's D value, and $N_s$ Number of sites.
