## Supplementary material for "Diversity and genetic structure within a Mexican maize race reveal consistent biocultural processes across geographic scales"

**Table S1. Sampling details table**

(see as a separate file)

### Supplementary Methods for Choosing Filtering Parameters

Filtering decisions were made following the pipeline described in "Speciation & Population Genomics: a how-to-guide" (https://speciationgenomics.github.io/filtering_vcfs/) by J. Meier and M. Ravinet. Analyses were conducted to make filtering decisions, mean depth per site, and variant missingness per site. We explored different quality statistics using a subset of 100,000 and 100 SNP based on which the parameters were decided to filter the final set. The quality analysis revealed that most sites had high Phred scores, indicating reliable calls, and thus no additional quality filtering was needed. The mean read depth per site was low but acceptable, ranging from 0.70 to 1.98, sufficient for maize. An 80% threshold for missing data was set, allowing for up to 20% data absence per site. Consequently, filtering was based on read depth and the proportion of missing data.

*Site or variant quality.* The probability distribution was obtained, as well as descriptive statistics values for the quality of sequences in all SNP, 4 959 703 (Fig. S1 and S2), 100 000 (Fig. S3), and 100 (Fig. S4) randomly selected variants. All variants were used for consistency in the mean and median of the randomly selected variants for the following analyses. Consistency in the value of the data quality was provided by the mode (Table S2). The quality scores observed were very high for all sites; recall that a score phred of 30 phred represents the probability that one variant in a thousand (1/1000 SNP) has been miscalled. Most sites exceeded this, suggesting that there were many high-confidence calls (min 30 for all variants and random). This may suggest a sufficient reading depth (high likelihood). It was concluded that most of the sites were of high quality, so there was no need for filtering. This coincided with the quality analysis performed using FastQC.

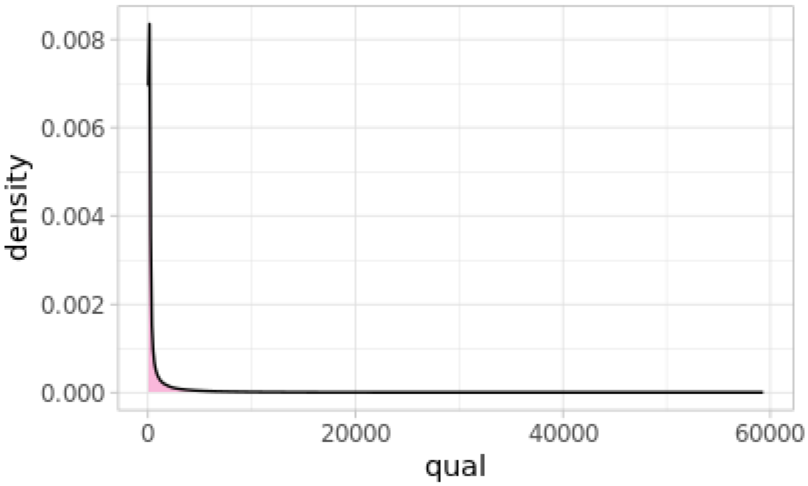


**Fig. S1 Site or variant quality.** Distribution of site quality for all SNP (4,959,703).

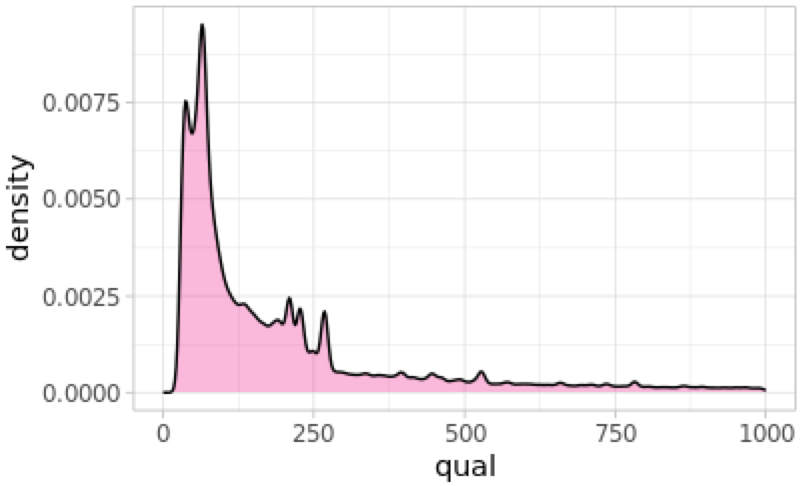


**Fig. S2 Site or variant quality.** Distribution of site quality for all SNP (4,959,703) xlim (0,1000).

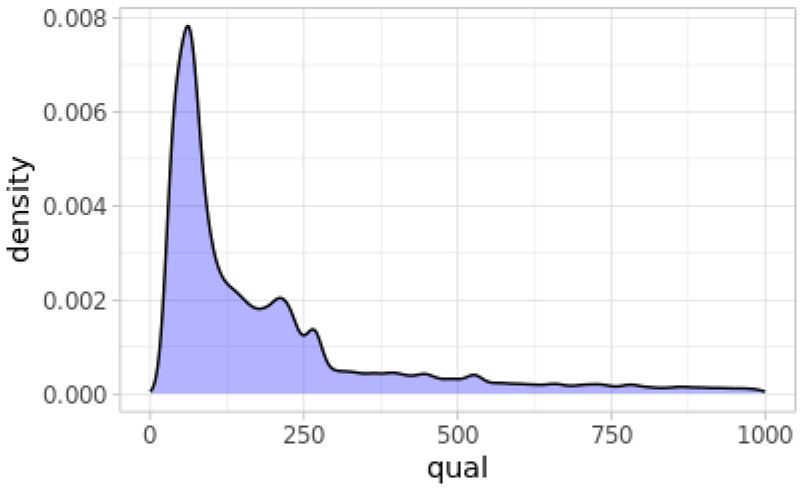


**Fig. S3 Site or variant quality.** Distribution of site quality for 100,000 SNP, xlim (0,1000).

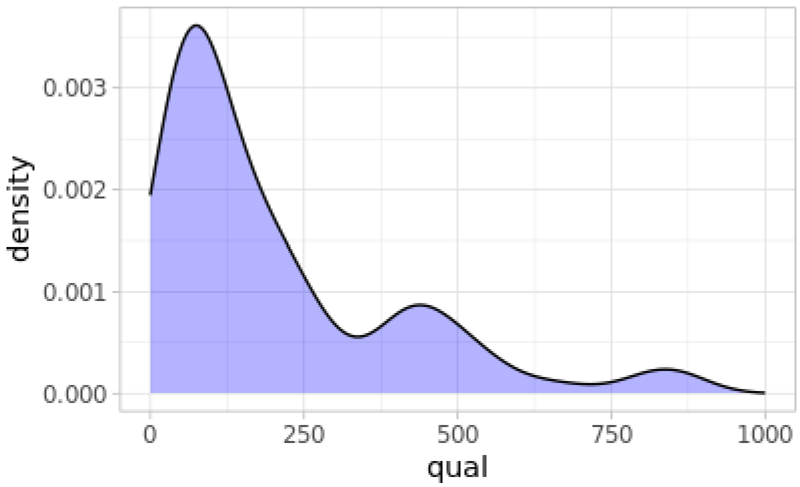


**Fig. S4 Site or variant quality.** Distribution of site quality for 100 SNP, xlim (0,1000).

**Table S2.** Descriptive statistics values for the quality of sequences for three subsets of the data (all, 100,000, and 100 SNP)

| metric | 4 959 703 SNP | 100 000 SNP | 100 SNP |
|---|---|---|---|
| minimum | 30 | 30 | 30 |
| maximum | 59 305.10 | 40 192.80 | 15 315.40 |
| median | 156.77 | 156.86 | 177.65 |
| mean | 888.74 | 883.66 | 893.23 |

*Mean depth per site or variant mean depth.* Very few variants with an extremely high coverage were observed. The median was 0.7083 and the mean was 0.9942, min = 0; max = 11.0156, (Fig. S5).

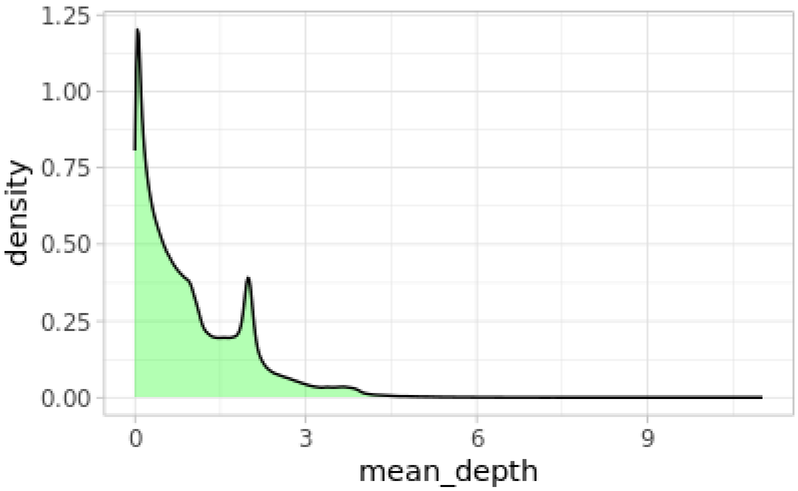


**Fig. S5 Mean depth per site excludes extreme values.** Distribution of mean depth per site for all SNP (4,959,703).

To determine the minimum and maximum depths, the median and mean should be considered, which in our case are 0.70 and 0.99, respectively, indicating that we had very few readings per site for all individuals. The minimum depth was 0.70, and the maximum depth would be the result of multiplying the mean by two (0.99 x 2 = 1.98). Therefore, the range of choice was from 0.70 to 1.98. For filtering, 2x (min) and 4x (max) were considered because readings higher than 2x are available (Fig. S6). This range of depth is sufficient for maize as shown by previous studies (Rojas-Barrera et al. 2019), due to the high quality of the reference genome.

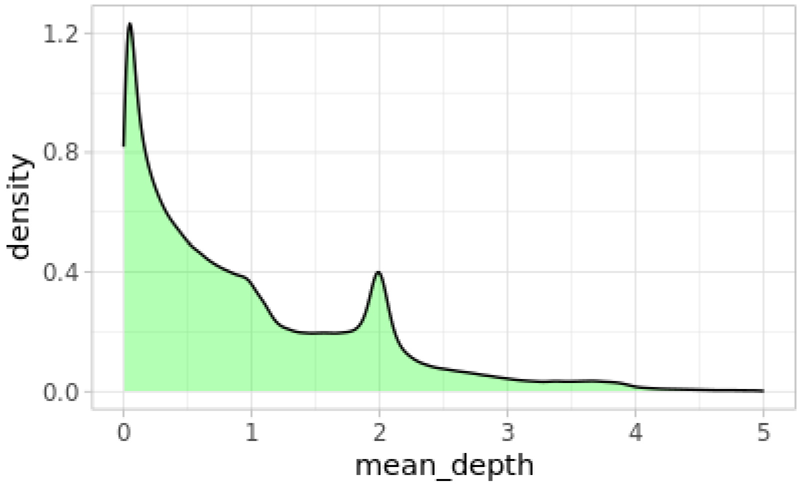


**Fig. S6 Mean depth per site**. Distribution of mean depth per site for all SNP (4,959,703).

*Variant Missingness per site.* There are numerous individuals with a high percentage of missing data (greater than 0.75), and there is also a significant number of individuals with relatively few missing data (less than 0.40) (Fig. S7). Typically, missingness of 0.75-0.95 is considered. In our case, we will use 0.80 (--max-missing 0.80): a site can have a maximum of 80% or 20% missing data. In other words, there may be missing data in 80% of the individuals.

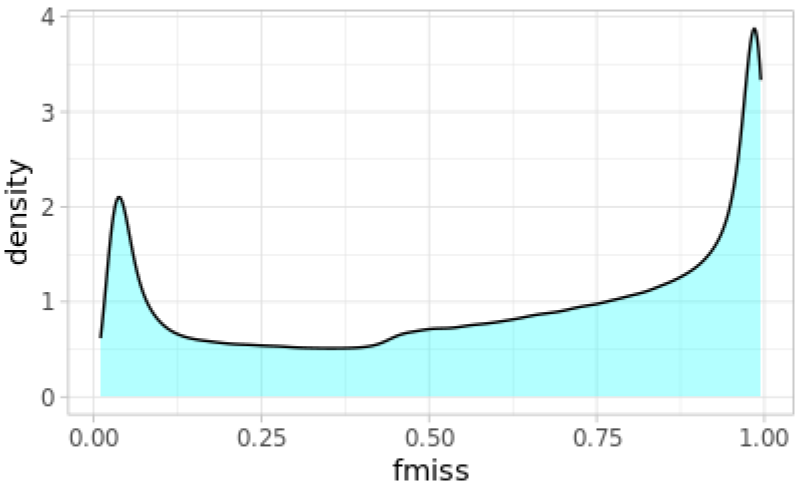


**Fig. S7 Variant Missingness per site.** Distribution of variant missingness per site for all SNP (4 959 703). The X-axis ranges from 0 to 1, where 0 means that the threshold means we will tolerate a 100% call-rate (there is no missing data for the variant in question) and 1 means 0% call-rate (the variant is completely missing in all the samples).

**Supplementary for CVE's plot**

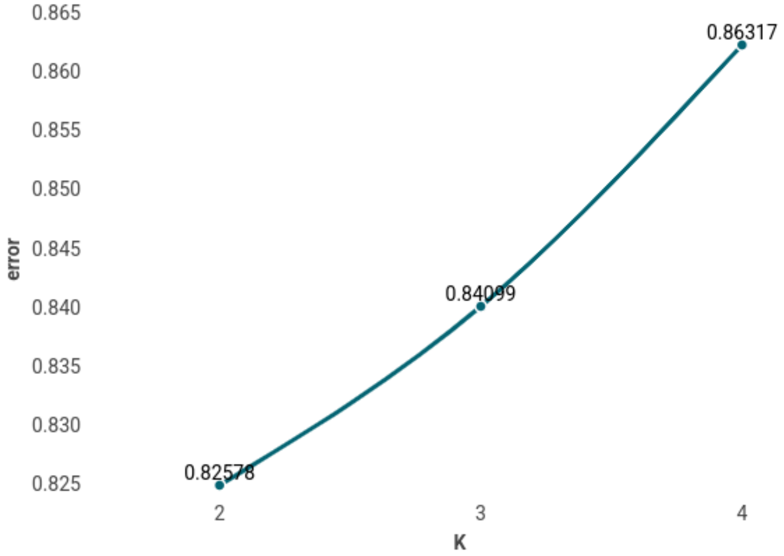


**Fig. S8 Plot of ADMIXTURE cross validation error from K=2 through K=4**. We chose K=2 to analyze the SNP data, as the value that minimizes the error. The value for the cross-validation error for each run (1-10) for the different K's was the same.

### Supplementary Plot for population genetics statistical tests

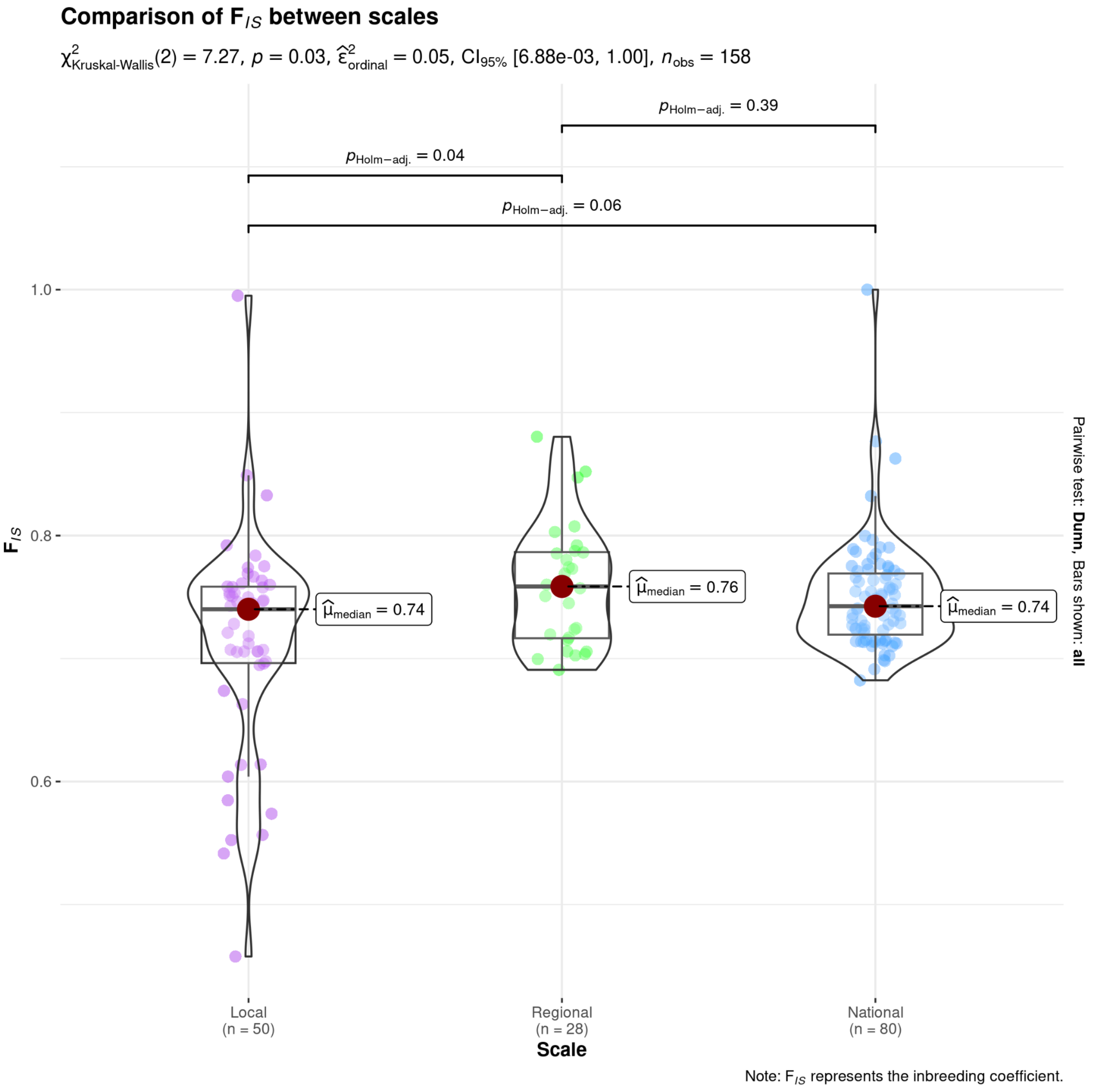


**Fig. S9. Kruskal-Wallis and Dunn test for $F_{IS}$.** Significant differences were only found between Local and Regional ($p$ = 0.04, $p$ < 0.05).

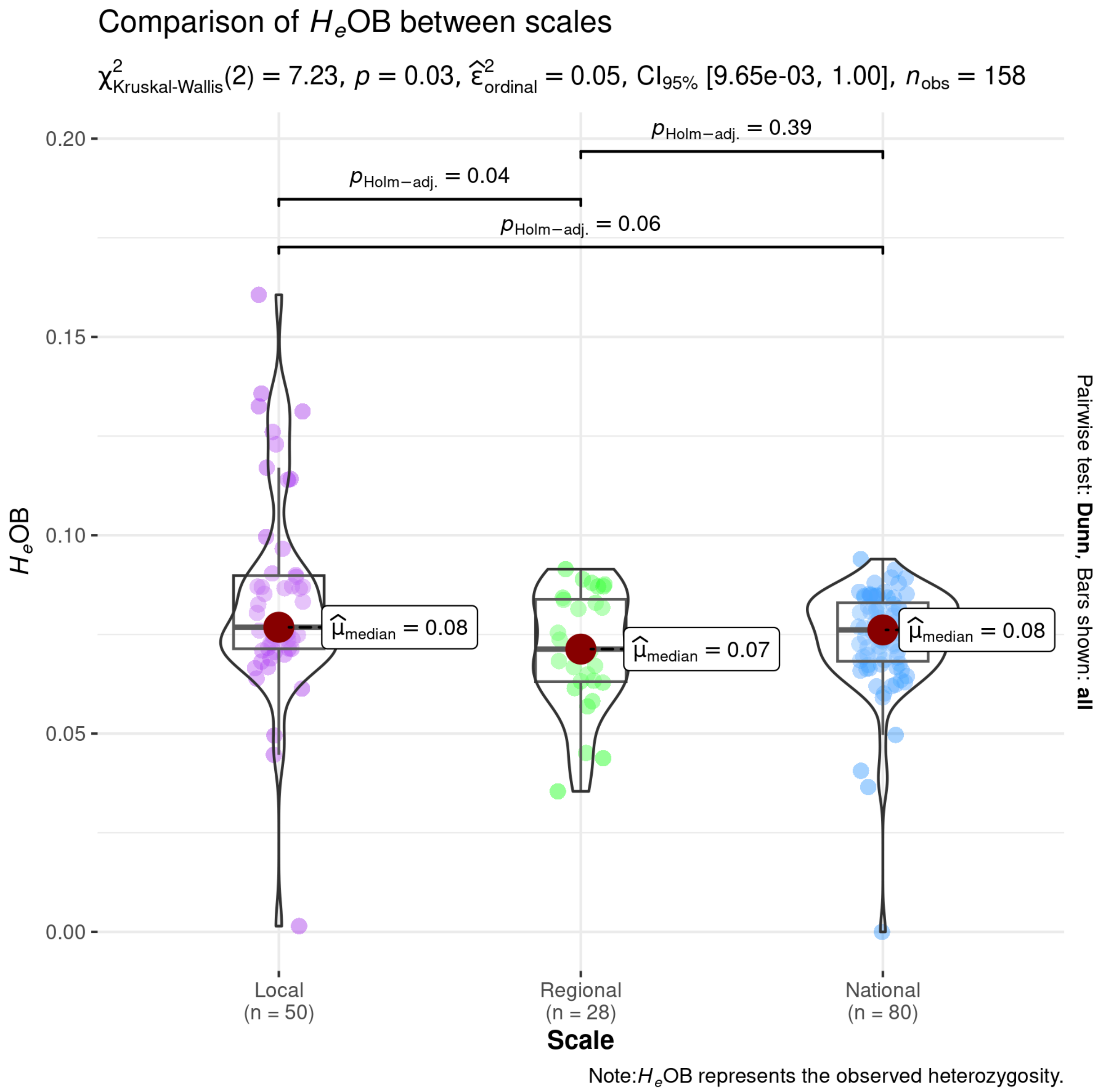


**Fig. S10. Kruskal-Wallis and Dunn test for $H_e$OB.** Significant differences were only found between Local and Regional ($p$ = 0.04, $p$ < 0.05).

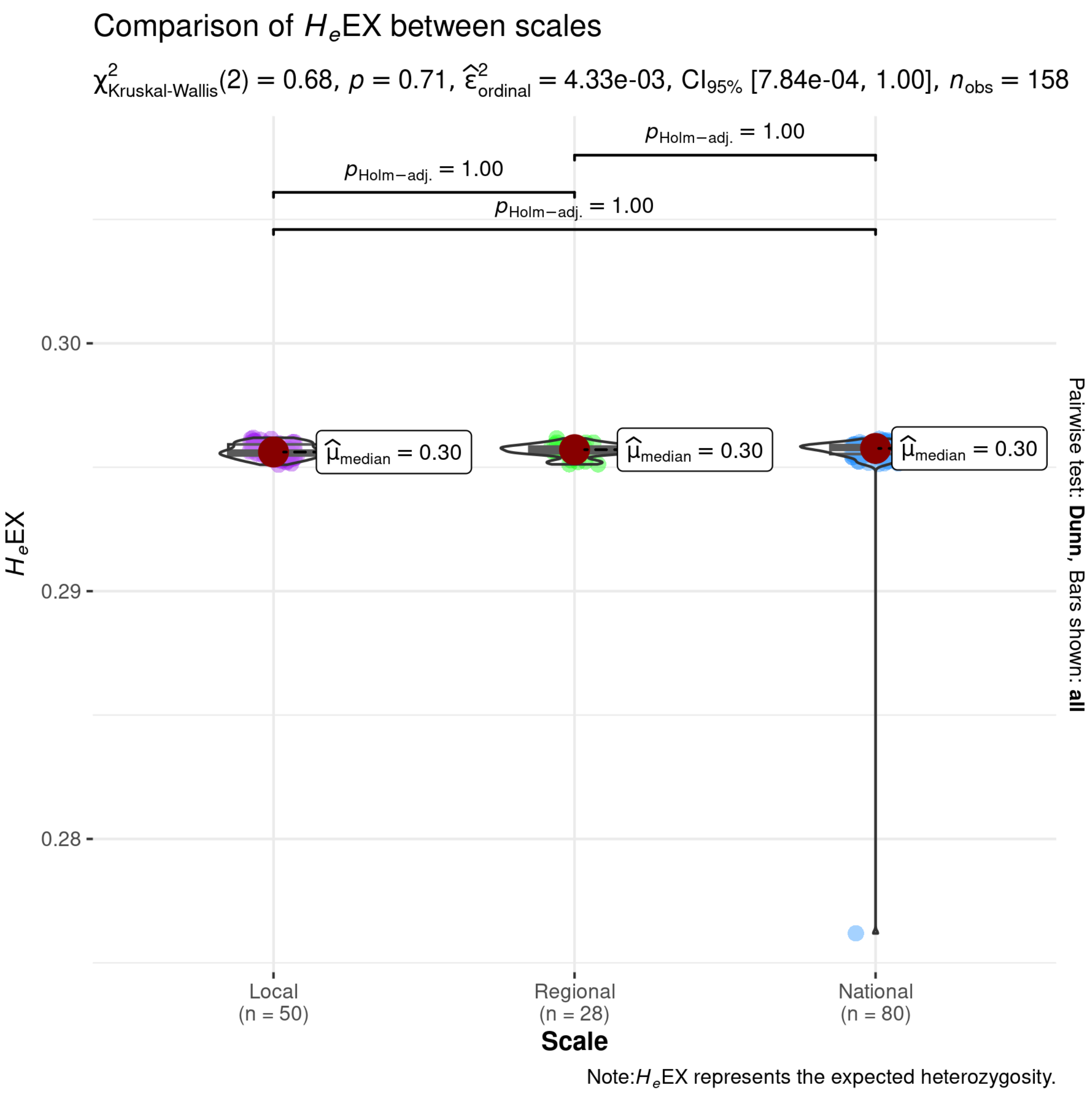


**Fig. S11. Kruskal-Wallis and Dunn test for $H_e$EX.** No significant differences were found.

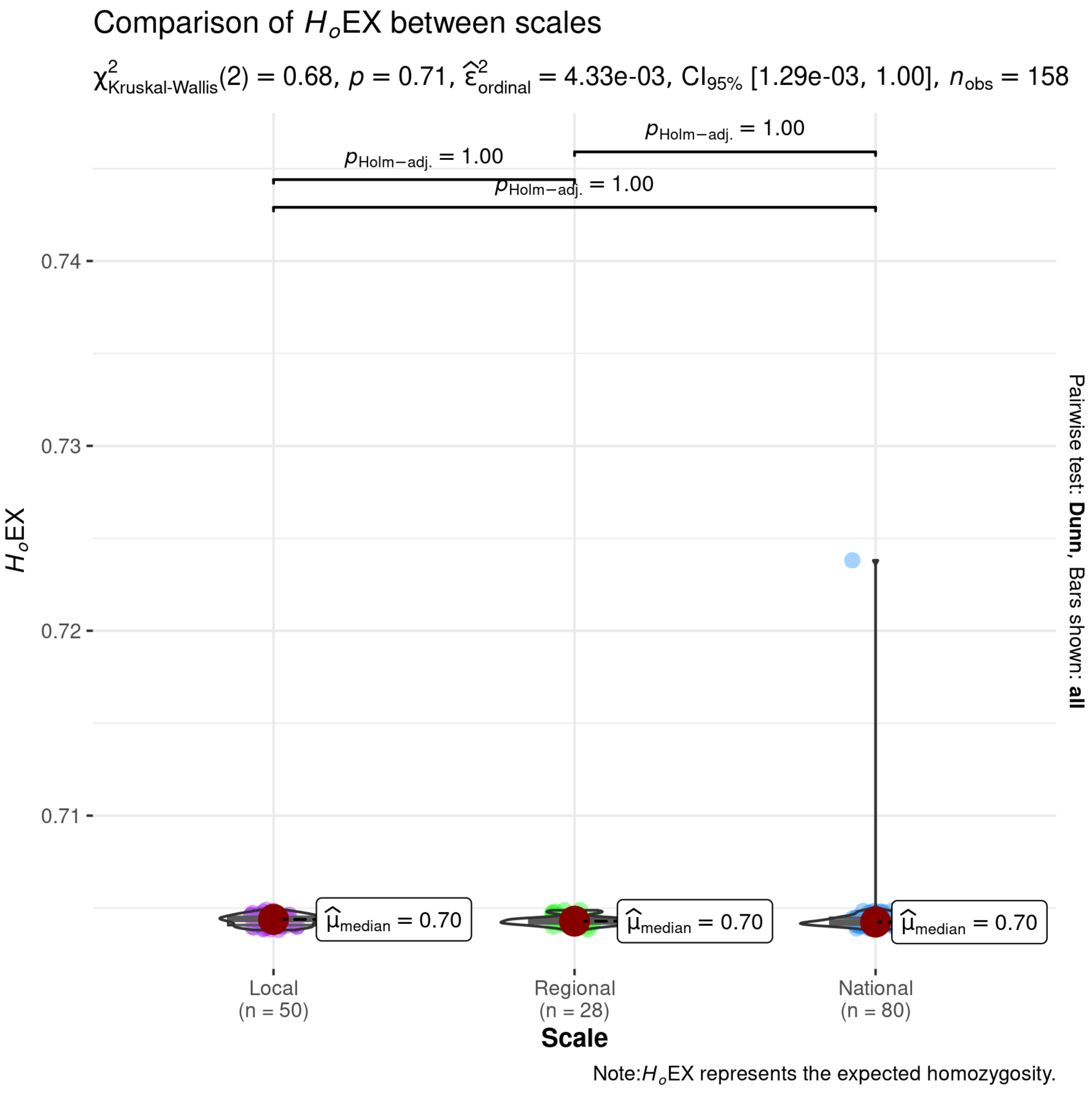


**Fig. S12. Kruskal-Wallis and Dunn test for $H_o$EX.** No significant differences were found.

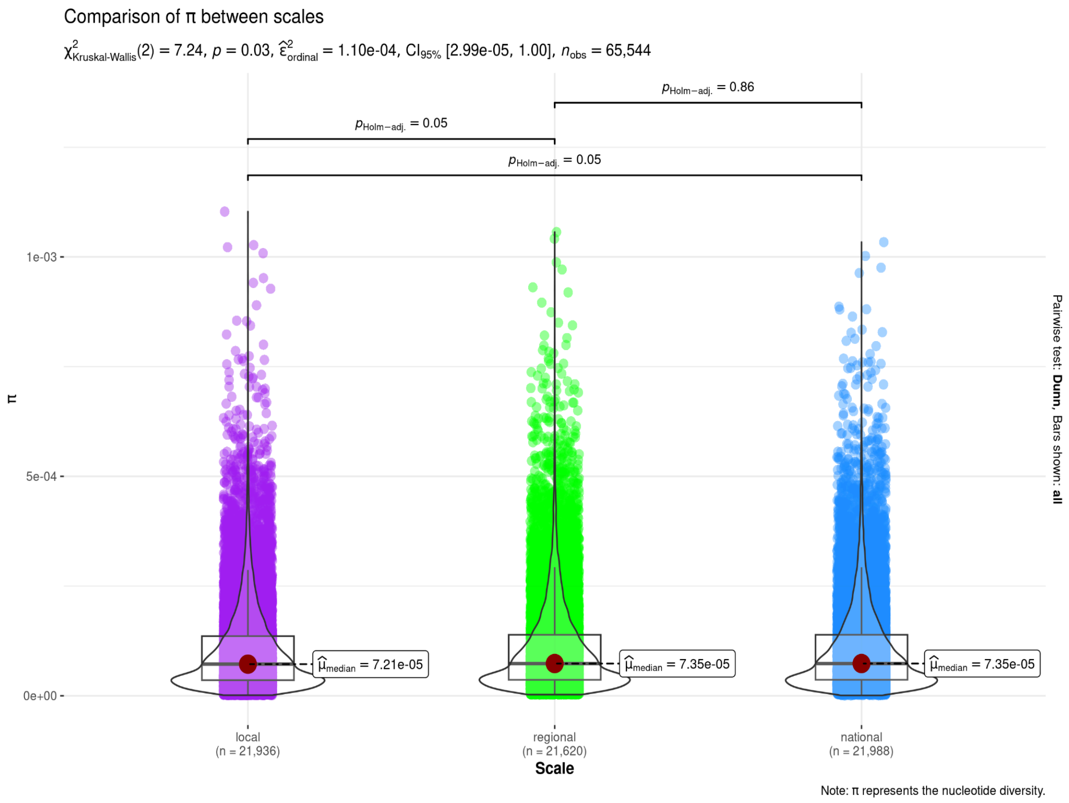


**Fig. S13. Kruskal-Wallis and Dunn test for π.** No significant differences were found.

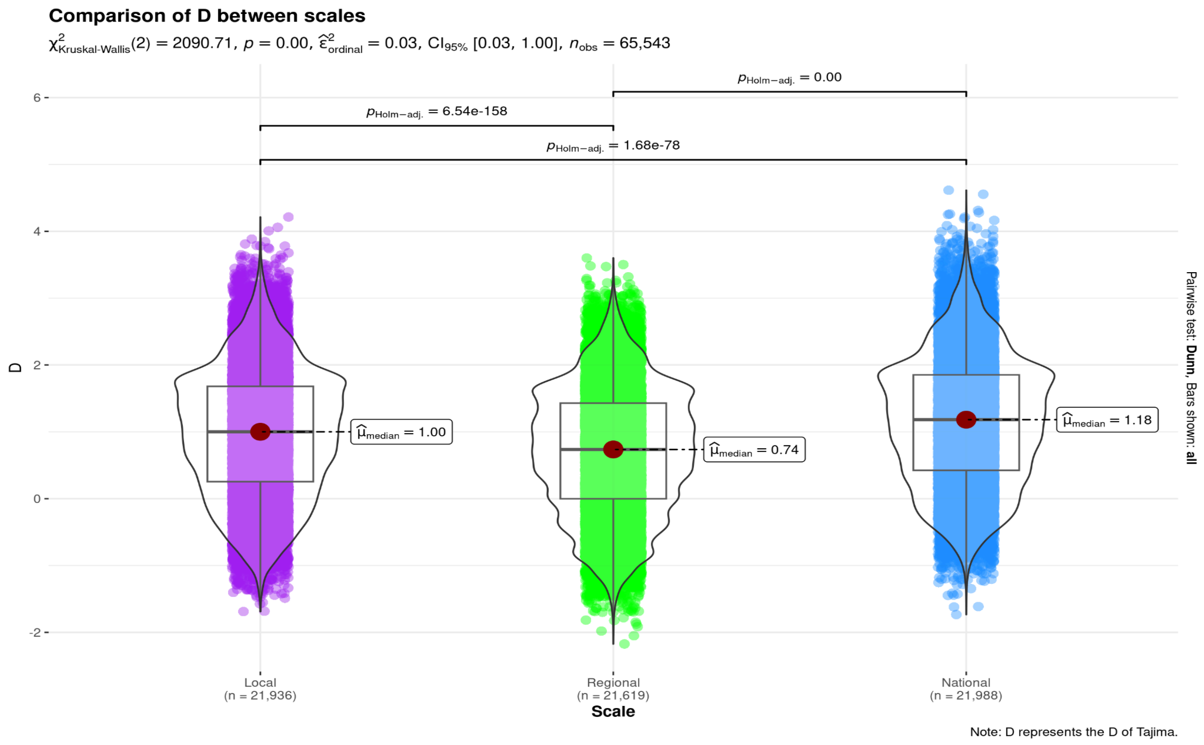


**Fig. S14. Kruskal-Wallis and Dunn test for *D*.** Significant differences were found between all groups.

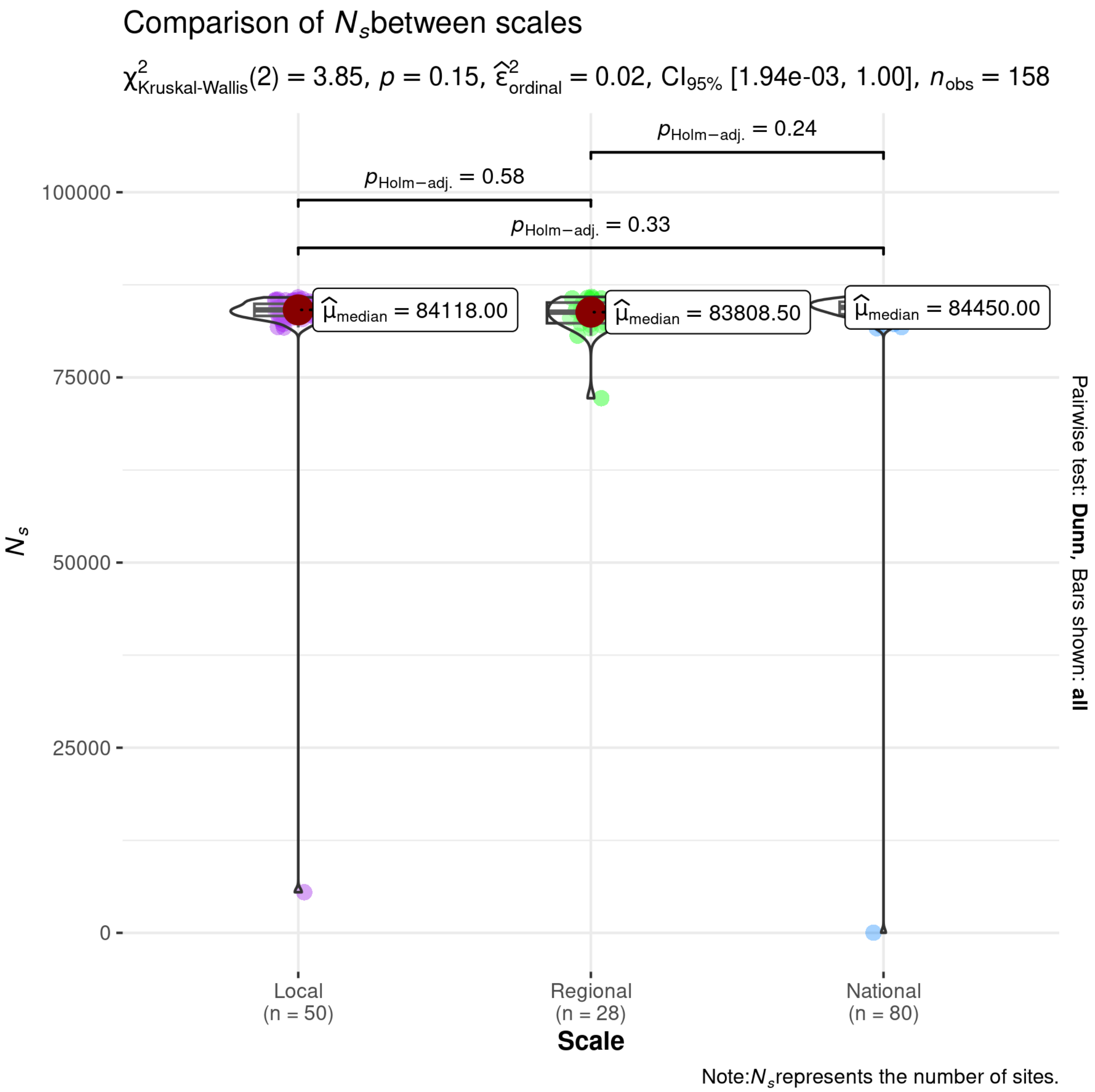


**Fig. S15. Kruskal-Wallis and Dunn test for $N_s$.** No significant differences were found.

**Table S3. Mean and standard deviation (sd) of genetic diversity summary statistics* for each accession**

| accession ID | $F_{IS}$ mean | $F_{IS}$ sd | $H_e$OB mean | $H_e$OB sd | $H_e$EX mean | $H_e$EX sd | $H_O$EX mean | $H_O$EX sd | *n* |
|---|---|---|---|---|---|---|---|---|---|
| CHIS_E52 | 0.73 | 0.021 | 0.080 | 0.006 | 0.296 | 0 | 0.704 | 0 | 2 |
| CHIS_E59 | 0.778 | 0.012 | 0.066 | 0.004 | 0.296 | 0 | 0.704 | 0 | 2 |
| CHIS_L04 | 0.739 | 0.047 | 0.077 | 0.014 | 0.296 | 0 | 0.704 | 0 | 4 |
| CHIS_L08 | 0.76 | 0.053 | 0.071 | 0.016 | 0.296 | 0 | 0.704 | 0 | 4 |
| CHIS_L33 | 0.815 | 0.12 | 0.055 | 0.035 | 0.296 | 0 | 0.704 | 0 | 4 |
| CHIS_L34 | 0.74 | 0.032 | 0.077 | 0.01 | 0.296 | 0 | 0.704 | 0 | 4 |
| CHIS_L35 | 0.719 | 0.02 | 0.083 | 0.006 | 0.296 | 0 | 0.704 | 0 | 4 |
| CHIS_L36 | 0.679 | 0.142 | 0.095 | 0.042 | 0.296 | 0 | 0.704 | 0 | 4 |
| CHIS_L37 | 0.673 | 0.148 | 0.097 | 0.044 | 0.296 | 0 | 0.704 | 0 | 4 |
| CHIS_L41 | 0.708 | 0.094 | 0.086 | 0.028 | 0.296 | 0 | 0.704 | 0 | 4 |
| CHIS_L42 | 0.749 | 0.022 | 0.074 | 0.006 | 0.296 | 0 | 0.704 | 0 | 4 |
| CHIS_L43 | 0.601 | 0.041 | 0.118 | 0.012 | 0.296 | 0 | 0.704 | 0 | 6 |
| CHIS_L44 | 0.721 | 0.039 | 0.083 | 0.012 | 0.296 | 0 | 0.704 | 0 | 4 |
| CHIS_L45 | 0.735 | 0.023 | 0.078 | 0.007 | 0.296 | 0 | 0.704 | 0 | 4 |
| CHIS_R02 | 0.757 | 0.026 | 0.072 | 0.008 | 0.296 | 0 | 0.704 | 0 | 4 |
| CHIS_R05 | 0.771 | 0.08 | 0.068 | 0.024 | 0.296 | 0 | 0.704 | 0 | 4 |
| CHIS_R06 | 0.764 | 0.065 | 0.07 | 0.019 | 0.296 | 0 | 0.704 | 0 | 4 |
| CHIS_R10 | 0.773 | 0.04 | 0.067 | 0.012 | 0.296 | 0 | 0.704 | 0 | 4 |
| CHIS_R11 | 0.763 | 0.074 | 0.070 | 0.022 | 0.296 | 0 | 0.704 | 0 | 4 |
| CHIS_R40 | 0.733 | 0.043 | 0.079 | 0.013 | 0.296 | 0 | 0.704 | 0 | 4 |
| GRO_N13 | 0.775 | 0.054 | 0.066 | 0.016 | 0.296 | 0 | 0.704 | 0 | 4 |
| GRO_N14 | 0.753 | 0.031 | 0.073 | 0.009 | 0.296 | 0 | 0.704 | 0 | 4 |
| GRO_N15 | 0.754 | 0.033 | 0.073 | 0.01 | 0.296 | 0 | 0.704 | 0 | 4 |
| GRO_N16 | 0.746 | 0.032 | 0.075 | 0.01 | 0.296 | 0 | 0.704 | 0 | 4 |

| | | | | | | | | | |
|---|---|---|---|---|---|---|---|---|---|
| GRO_N18 | 0.776 | 0.019 | 0.066 | 0.006 | 0.296 | 0 | 0.704 | 0 | 4 |
| GRO_N19 | 0.737 | 0.037 | 0.078 | 0.011 | 0.296 | 0 | 0.704 | 0 | 4 |
| GRO_N20 | 0.733 | 0.018 | 0.079 | 0.005 | 0.296 | 0 | 0.704 | 0 | 3 |
| GRO_N22 | 0.744 | 0.018 | 0.076 | 0.005 | 0.296 | 0 | 0.704 | 0 | 4 |
| GRO_N23 | 0.8 | 0.135 | 0.059 | 0.04 | 0.291 | 0.01 | 0.709 | 0.01 | 4 |
| GRO_N24 | 0.766 | 0.074 | 0.069 | 0.022 | 0.296 | 0 | 0.704 | 0 | 4 |
| GRO_N26 | 0.723 | 0.011 | 0.082 | 0.003 | 0.296 | 0 | 0.704 | 0 | 4 |
| HGO_E50 | 0.736 | 0.037 | 0.078 | 0.011 | 0.296 | 0 | 0.704 | 0 | 2 |
| NAY_N27 | 0.724 | 0.013 | 0.082 | 0.004 | 0.296 | 0 | 0.704 | 0 | 4 |
| NAY_N29 | 0.76 | 0.069 | 0.071 | 0.021 | 0.296 | 0 | 0.704 | 0 | 4 |
| NAY_N30 | 0.719 | 0.034 | 0.083 | 0.01 | 0.296 | 0 | 0.704 | 0 | 4 |
| NAY_N47 | 0.74 | 0.041 | 0.077 | 0.012 | 0.296 | 0 | 0.704 | 0 | 4 |
| OAX_N31 | 0.737 | 0.019 | 0.078 | 0.006 | 0.296 | 0 | 0.704 | 0 | 4 |
| OAX_N32 | 0.77 | 0.022 | 0.068 | 0.007 | 0.296 | 0 | 0.704 | 0 | 5 |
| SLP_E49 | 0.706 | 0.01 | 0.087 | 0.003 | 0.296 | 0 | 0.704 | 0 | 2 |
| SLP_E51 | 0.741 | 0.024 | 0.077 | 0.007 | 0.296 | 0 | 0.704 | 0 | 2 |
| VER_E48 | 0.733 | 0.027 | 0.079 | 0.008 | 0.296 | 0 | 0.704 | 0 | 2 |
| VER_E56 | 0.772 | 0.005 | 0.067 | 0.002 | 0.295 | 0 | 0.705 | 0 | 2 |
| VER_E57 | 0.769 | 0.012 | 0.068 | 0.003 | 0.295 | 0 | 0.705 | 0 | 2 |

* $F_{IS}$ Inbreeding coefficient, $H_e$OB Observed heterozygosity, $H_e$Ex Expected heterozygosity, $H_o$Ex Expected homozygosity, *n* number of samples per accession.
